# Motor cortex can directly drive the globus pallidal neurons in a projection neuron type dependent manner in rat

**DOI:** 10.1101/677377

**Authors:** Fuyuki Karube, Susumu Takahashi, Kenta Kobayashi, Fumino Fujiyama

## Abstract

The basal ganglia (BG) are critical for the control of motor behaviors and for reinforcement learning. Here, we demonstrate in rats that primary and secondary motor areas (M1 and M2) make functional synaptic connections in the globus pallidus (GP), not usually thought of as an input site of the BG using optogenetics and morphological analysis. The cortical excitation in the GP was as strong as that in the STN. GP neurons projecting to the striatum were preferentially innervated by the motor cortex. Morphological observation revealed that the density of axonal boutons from motor cortices in the GP was approximately 30% of that in the striatum, but was comparable to that in the STN. M1 and M2 projected differentially to the BG in terms of topography and substructures. These results suggest that cortico-pallidal innervation is an additional excitatory input to the BG, and can affects behaviors via the cortex-basal ganglia-thalamus loop.

## Introduction

Parallel loops of neural connections among the cerebral cortex, basal ganglia, and thalamus contribute to multiple aspects of behavior (Alexander, DeLong, & Strick, 1986; Nambu, 2008; Wei & Wang, 2016). The functions mediated by these loops depend on relevant cortical areas and brain regions receiving outputs of the basal ganglia (Hikosaka, 2007; Middleton & Strick, 2000). The loop containing the motor cortex is crucial for appropriate motor control, action selection, and movement-related learning. Dysfunction of the motor loop involves movement disorders such as Parkinsonian disease (Albin, Young, & Penney, 1989; DeLong, 1990; Middleton & Strick, 2002; Nambu, 2008; Nambu et al., 2000; Parent & Hazrati, 1995a; Redgrave et al., 2010; Wichmann & DeLong, 1996). Cortical projections drive three pathways in the basal ganglia: the direct, the indirect, and the hyperdirect pathways (Bolam, Hanley, Booth, & Bevan, 2000; Y. Smith, Bevan, Shink, & Bolam, 1998). The direct and indirect pathways are mediated by two distinct types of striatal medium spiny neurons (MSNs), termed the direct- and indirect-pathway MSNs (dMSNs and iMSNs). dMSNs project to the output nuclei of the basal ganglia, namely the substantia nigra (SN) pars reticulata (SNr), and the globus pallidus internal segment, the latter termed the entopeduncular nucleus (EP) in rodents. The iMSNs project to the globus pallidus external segment (GP in rodents), which interconnects with the subthalamic nucleus (STN). In turn, both STN and GP also innervate the output nuclei. The hyperdirect pathway involves direct cortical projections to the STN (Nambu, Tokuno, & Takada, 2002), which provides the fastest information flow among the three pathways (Nambu et al., 2000). The behavioral functions of these pathways are gradually being elucidated, although recent findings propose refinement and reappraisal of the classical views of the functional roles of distinct MSNs (Calabresi, Picconi, Tozzi, Ghiglieri, & Di Filippo, 2014; Cui et al., 2013; Isomura et al., 2013; Vicente, Galvao-Ferreira, Tecuapetla, & Costa, 2016). According to the traditional model, the direct pathway promotes the execution of desired actions, whereas the indirect pathway prevents the execution of competing actions (Friend & Kravitz, 2014; Nambu, 2007; Vicente et al., 2016), and the hyperdirect pathway emergently cancels or switches imminent movements (Frank, Samanta, Moustafa, & Sherman, 2007; Isoda & Hikosaka, 2008; Nambu et al., 2002; Schmidt, Leventhal, Mallet, Chen, & Berke, 2013). In this regard, basal ganglia activity and function are likely modulated by the cerebral cortex.

The rodent motor cortical area is composed of primary and secondary motor cortices, M1 and M2, respectively. The rodent M1 codes and conducts movement signals (Barth, Jones, & Schallert, 1990; Brown & Teskey, 2014) via direct projections to the brainstem and spinal cord, analogously to the primate M1. It has been debated whether the rodent M2, which has various other names such as the medial agranular cortex and the medial precentral cortex (Ebbesen et al., 2018; Svoboda & Li, 2018), is a functional counterpart of the primate premotor and/or supplementary motor areas (Barthas & Kwan, 2017; Svoboda & Li, 2018). Recent studies have revealed the functional role and significance of the rodent M2 (Guo et al., 2014; Hira et al., 2014; Hira et al., 2013; Li, Chen, Guo, Gerfen, & Svoboda, 2015; Manita et al., 2015; D. Miyamoto et al., 2016; Murakami, Shteingart, Loewenstein, & Mainen, 2017; Murakami, Vicente, Costa, & Mainen, 2014; Saiki et al., 2014; Soma et al., 2017; Sul, Jo, Lee, & Jung, 2011). The M2 integrates motor signals and multimodal sensory/internal state information [for review see Barthas and Kwan (2017)]. The M2 is also involved in the preparatory and movement phases of behavior (Guo et al., 2014; Heindorf, Arber, & Keller, 2018; Li et al., 2015; Murakami et al., 2017; Murakami et al., 2014), and learning of motor tasks (Cao et al., 2015; Kawai et al., 2015). Other frontal cortical areas are also related to motor behaviors, such as the orbitofrontal cortex and the cingulate motor area (Bissonette, Powell, & Roesch, 2013; Friedman et al., 2015; Nakayama, Yokoyama, & Hoshi, 2015; Passingham & Wise, 2012; Paus, 2001; Schoenbaum, Roesch, Stalnaker, & Takahashi, 2009; Sul, Kim, Huh, Lee, & Jung, 2010). The neural circuitry underlying these functional roles and its effects on the basal ganglia and other subcortical regions are not fully known.

Neuron-type diversity is an important factor to understand the neural circuitry. It has been established that distinct neural types other than striatal dMSNs and iMSNs are present in the GP (Abdi et al., 2015; Abrahao & Lovinger, 2018; Cooper & Stanford, 2000; Dodson et al., 2015; Gittis et al., 2014; Hegeman, Hong, Hernandez, & Chan, 2016; Hernandez et al., 2015; H. Kita, 2007; Mallet et al., 2012; Mastro, Bouchard, Holt, & Gittis, 2014), EP (Y. Miyamoto & Fukuda, 2015; Wallace et al., 2017), STN (Baufreton et al., 2003; Xiao et al., 2015), and SN (Kim, Ghazizadeh, & Hikosaka, 2014; Lerner et al., 2015; Matsumoto & Hikosaka, 2009; Menegas et al., 2015). These neuronal populations differ in molecular profile, projection targets, and synaptic inputs. In addition, each nucleus of the basal ganglia contains functional/morphological subdomains (Crittenden & Graybiel, 2011; Fujiyama, Takahashi, & Karube, 2015; Hontanilla, Parent, de las Heras, & Gimenez-Amaya, 1998; H. Kita & Kita, 2001; Parent, Fortin, Cote, & Cicchetti, 1996) correlated with the distribution of glutamatergic innervation (Crittenden & Graybiel, 2011; Eblen & Graybiel, 1995; Fujiyama, Unzai, Nakamura, Nomura, & Kaneko, 2006; Kincaid & Wilson, 1996; J. B. Smith et al., 2016). Even if innervated by the same set of cortical inputs, each cell type can behave differently, recruit distinct neural circuitry, and lead to various behavioral outputs. Therefore, any account of the functionality of cortical projections to the basal ganglia must consider cell type and subregion.

Cortico-basal ganglia projections are topographically and somatotopically organized (Crittenden & Graybiel, 2011; Gabbott, Warner, Jays, Salway, & Busby, 2005; Nambu, 2011; Shipp, 2016; Voorn, Vanderschuren, Groenewegen, Robbins, & Pennartz, 2004). In the rodent, frontal cortical areas project to the dorsal striatum (Barth et al., 1990; Ebrahimi, Pochet, & Roger, 1992; Reep & Corwin, 1999; Reep et al., 2008; Rouiller, Moret, & Liang, 1993). Recently, topographical maps of cortico-striatal projections have been described in detail (Hintiryan et al., 2016; Hunnicutt et al., 2016; Mailly, Aliane, Groenewegen, Haber, & Deniau, 2013). Cortical projections to the striatum are dependent on the neuron type of origin (Hooks et al., 2018) and individual cortical neurons generally project to multiple sites (T. Kita & Kita, 2012; Shepherd, 2013; Shibata, Tanaka, Hioki, & Furuta, 2018; but see also. Y. Smith, Wichmann, & DeLong, 2014). Thus, basal ganglia nuclei innervated by the same or different cortical areas can be affected simultaneously, in turn, their activities are modulated via the intra-basal ganglia circuitry (Bogacz, Martin Moraud, Abdi, Magill, & Baufreton, 2016; Wei & Wang, 2016). The aim of the present study was to investigate cortical projections to multiple nuclei in the basal ganglia from two motor areas, M1 and M2, in a subregion- and cell type-specific manner. Using a combination of neural tracing with immunofluorescence, we demonstrate that M1 and M2 project to different subregions of each basal ganglia nucleus, and that in the GP, cortical axon collaterals and boutons have topographic distributions that depend on the cortical area of origin (Naito & Kita, 1994). Using morphological and electrophysiological experiments combined with optogenetics, we demonstrate that cortico-pallidal synapses are effective and specific for post-synaptic GP cell types. We discuss the potential roles of cortico-pallidal projections in relation to other basal ganglia nuclei.

## Results

### Motor cortex innervates the GP

We observed cortical axon collaterals in the GP using conventional tracers (BDA or PHA-L; Fig. 1A, 1C; Fig. 1-figure supplement. 1) or adenoassociated virus (AAV) vectors (Fig. 2A, 5A; Fig. 1-figure supplement 2), as reported previously (Naito & Kita, 1994).

**Fig. 1.**
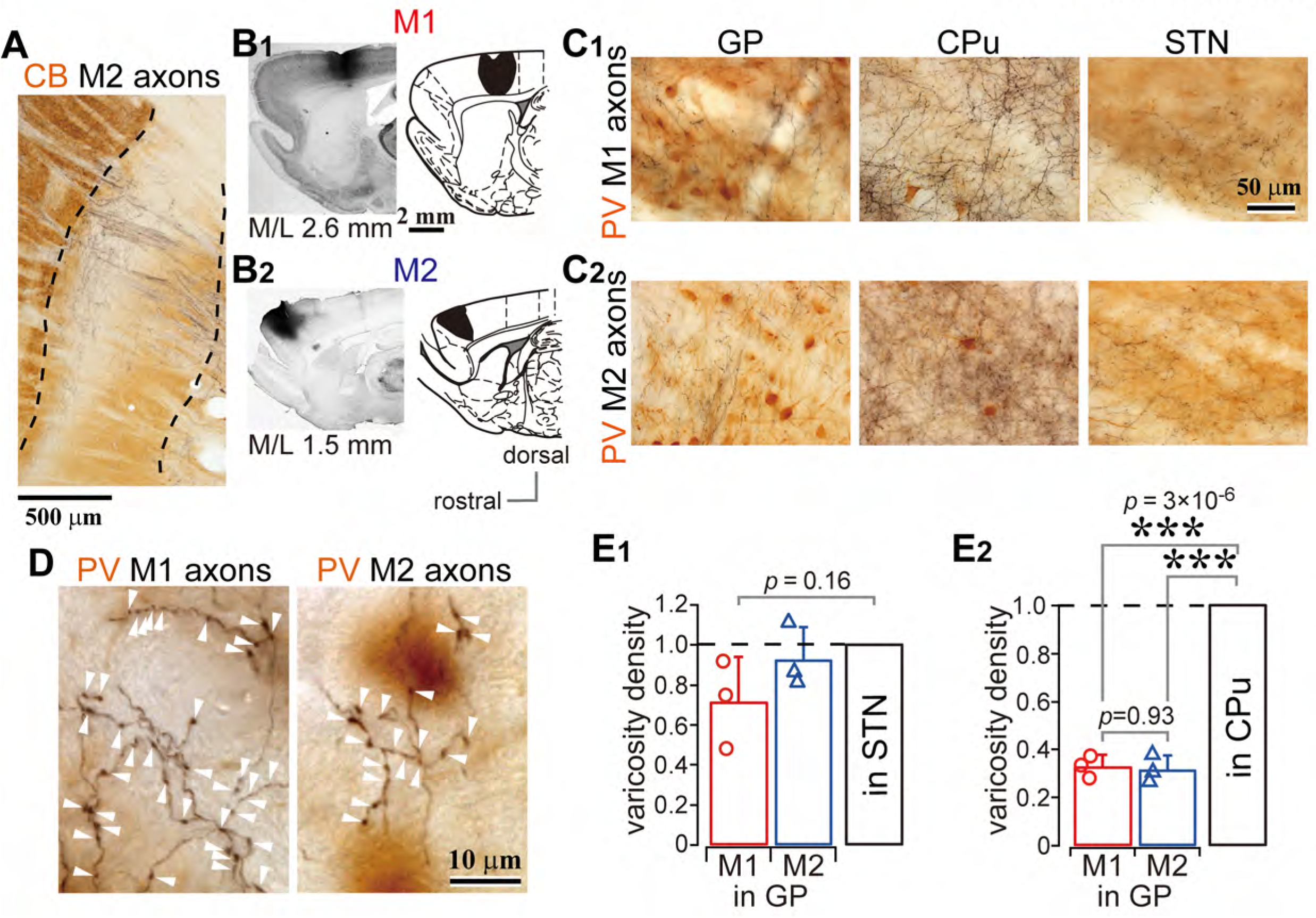
Motor cortical axons project to the globus pallidus (GP). (A) A representative image of axons in the GP originated from the secondary motor cortex (M2), labeled with biotinylated dextran amine (BDA; visualized in black). Axon collaterals in the GP are predominantly distributed in the calbindin (CB)-negative subregions. CB is visualized in brown. (B) Images and drawings of BDA injection sites into the primary and secondary motor areas (M1 and M2, respectively). Tracer was deposited across the entire thickness of the cortex. (C) Images of axon distributions in the GP, striatum, and subthalamic nucleus (STN) from M1 and M2. The sections were counter-stained with an anti-parvalbumin (PV) antibody, visualized in brown. (D) Magnified views of cortical axon varicosities (arrowheads) in the GP; **left**, M1 axons; **right**, M2 axons. The images are composites from multiple focal planes, and show that the PV neurons are unlikely to be contacted by axon varicosities. (E) Comparison of axon varicosity density in GP with that in the STN and striatum (*N* = 3 rats). The axon varicosity density in the GP as normalized values to that in the STN (E1), or to that in the striatum (E2).

**Fig. 2.**
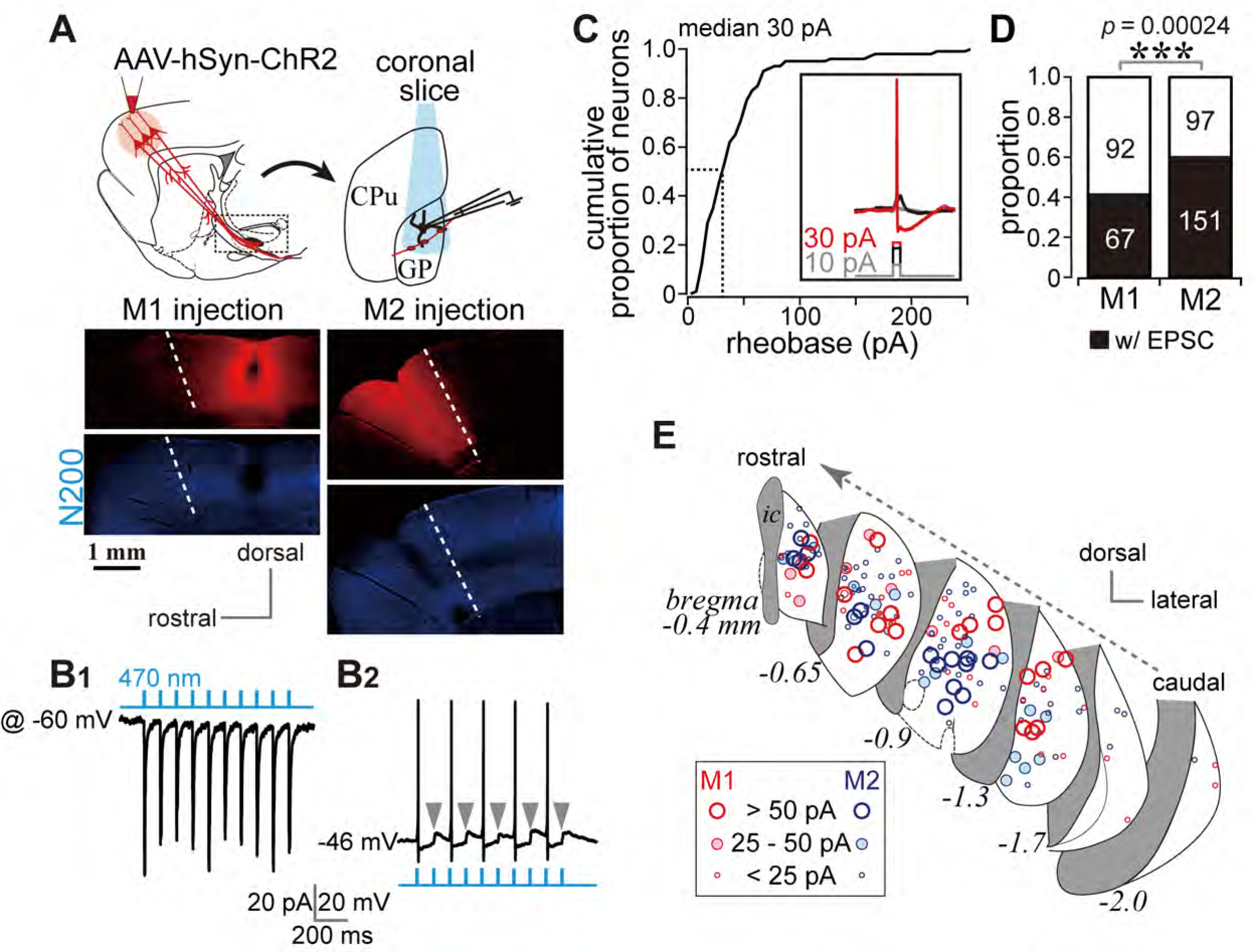
Photoactivation of motor cortical terminals evokes excitation in GP neurons. (A) Schematic (**top**) of AAV encoding channel rhodopsin 2 and mCherry injection into the motor cortex for *ex vivo* recordings using coronal slices. Examples of AAV injection sites are shown in the **middle** panels (red). Images of immunofluorescence for neurofilament 200 kDa (N200, **bottom**), used for identification of the M1/M2 border (white dotted lines). (B1) A representative voltage clamp trace (held at −60 mV) showing inward currents in GP neurons elicited by 5-ms blue light pulses (470 nm). (B2) A representative current clamp trace showing photoinduced action potentials and excitatory postsynaptic potentials (EPSPs, arrowheads). (C) Cumulative histogram of the rheobase current of GP neurons. Note that 25 to 30 pA is sufficient to elicit action potentials in half of GP neurons (*N* = 100). (D) Proportion of GP neurons innervated by M1 or M2 terminals. The number of neurons is shown in bars. M2 more frequently innervated the GP than did M1. (E) Location of GP neurons innervated by M1 (red circle) or M2 (blue circle). Note the topographic distribution of M1 and M2 innervation. The size of circles represents the amplitude of optically evoked currents,

Cortico-pallidal axon collaterals were exclusively ipsilateral (Fig. 1-figure supplement 2), implying that the layer 5 (L5) pyramidal tract (PT) type of neuron could be providing this input, thereby conveying a copy of cortical motor signals. Cortical axons were differentially distributed in the GP depending on the cortical area of origin. M1 and M2 axons were observed across the broad extent of the medio-lateral axis (range: M/L 2.4–3.7 mm; see also Fig. 1-figure supplement 1). Axon collaterals issued from the main axons and often elongated along the dorso-ventral axis (Fig. 1A, Fig. 1-figure supplement 1). Combining with immunofluorescent staining for calbindin D-28k (CB), motor cortical axons, especially from M1, were preferentially found in CB-negative [CB(-)] subregions of the GP (Fig. 1A, Fig. 1-figure supplement 1).

These axons frequently formed small sized axonal boutons (Fig. 1C, 1D). We compared the density of cortical axonal boutons in the GP, STN, and striatum. The number of boutons was greater in the striatum than in the GP or STN (Fig. 1-figure supplement 3). Since the size and efficacy of the tracer labeling were not uniform across animals, the bouton density in the GP was normalized to that in the striatum or STN for each injection. Using this normalization, the bouton density in the striatum was found to be significantly higher than in the GP (*p* = 2.9×10^−6^ for M1 and *p* = 2.6×10^−6^ for M2 using one-way ANOVA followed by Tukey test) or STN (*p* = 0.0018 for M1; *p* = 0.0003 for M2; see also Fig. 1-figure supplement 3). Bouton densities in the GP and STN were not significantly different (Fig. 1E; *p* = 0.155 using one-way ANOVA). GP bouton density did not significantly differ between M1 and M2 axons (*p* = 0.32).

These data suggested that M1 and M2 can affect the GP as strongly as they affect the STN. However, because bouton density does not directly indicate synaptic efficacy, we evaluated the electrophysiological features of cortico-pallidal projections.

### Cortico-GP terminals elicit monosynaptic EPSCs

To confirm that the observed cortico-pallidal innervation was electrophysiologically functional, we conducted whole-cell patch clamp recordings combined with optogenetic stimulation of cortical terminals using *in vitro* slice preparations. Channel rhodopsin 2 (ChR2) was introduced into cortical neurons using *AAV-hSyn-H134R-mCherry* injection into either M1 or M2 (Fig. 2A). In voltage clamp mode at a holding potential of −60 mV, stimulation with a brief light pulse (5 ms, 470 nm) elicited inward currents in GP neurons (Fig. 2B1). The response was stable over repetitive stimulation (10 pulses at 2–10 Hz; Fig. 2B1). In current clamp mode, photoactivation elicited action potentials, although the action potential probability was affected by the spontaneous oscillation of the membrane potential (Fig. 2B2). To confirm that the photoactivated current that elicited action potentials was within the physiological range, we measured the minimum current required to induce action potentials (rheobase current) in GP neurons using 5-ms depolarizing pulses (Fig. 2C, inset). In half of GP neurons, the rheobase was less than 30 pA, and most GP neurons could emit an action potential with less than 100 pA of depolarization (*N* = 100 neurons; Fig. 2C). A depolarized membrane potential and a high input resistance (Table 1) led to easy induction of action potentials by small excitation.

**Table 1.** Electrophysiological properties of globus pallidus (GP) neurons

|  | GP<sub>STN</sub> (*N*=108) | GP<sub>CPu</sub> (*N* = 65) | *p* value |
| --- | --- | --- | --- |
| Mean membrane potential (mV) | -46.63 ± 4.95<br>(-34.2– -61.53) | -46.36 ± 5.9<br>(-33.59– -60.53) | 0.682 |
| Input resistance (MΩ) | 232.35 ± 131.87<br>(33.1–887.45) | 329.97 ± 168.19<br>(32.8–828.3) | 6.8×10<sup>-05</sup> *** |
| Time constant (ms) | 12.94 ± 10.23<br>(2.25–79.75) | 21.92 ± 13.11<br>(2.42–55.67) | 2.1×10<sup>-07</sup> *** |
| Sag potential (mV) | 7.23 ± 4.52<br>(1.25–22.02) | 8.71 ± 7<br>(0.63–35.18) | 0.3432 |
| Spike frequency at 100 pA depolarization<br>(Hz) | 48.46 ± 22.94<br>(0–103) | 35.81 ± 19.6<br>(2–85) | 0.00041 *** |
| Spike frequency at 500 pA depolarization<br>(Hz) | 99.14 ± 66.87<br>(1–244) | 35.24 ± 39.24<br>(1–172) | 1.8×10<sup>-09</sup> *** |
| Maximum spike frequency (Hz) | 135.04 ± 65.42<br>(11–263) | 72.14 ± 36.12<br>(2–172) | 1.0×10<sup>-09</sup> *** |
| Current at maximum spike frequency (pA) | 578.63 ± 277.97<br>(50–1000) | 345.24 ± 219<br>(50–1000) | 6.9×10<sup>-08</sup> *** |
| Spike height (mV) | $74.06 \pm 9.77$<br>(47.04–96.98) | $76.55 \pm 11.69$<br>(38.84–95.97) | 0.051 |
| Spike width (ms) | $0.95 \pm 0.29$<br>(0.63–2.29) | $1.19 \pm 0.32$<br>0.7–2.17 | $1.6 \times 10^{-09}$ *** |
| Spike threshold (mV) | $-37.57 \pm 4.79$<br>(-24.09– -48.11) | $-37.7 \pm 5.32$<br>(-23.14– -46.03) | 0.5351 |
| fAHP amplitude (mV) | $21.16 \pm 5.14$<br>(9.1–38.72) | $18.09 \pm 4.57$<br>(11.88–36.34) | $1.3 \times 10^{-05}$ *** |
| fAHP delay after spike peak (ms) | $0.98 \pm 0.39$<br>(0.55–2.7) | $1.31 \pm 0.55$<br>(0.6–3.8) | $3.4 \times 10^{-07}$ *** |
| sAHP amplitude (mV) | $18.26 \pm 4.2$<br>(9.47–32.87) | $16.54 \pm 4.71$<br>(10.38–29.78) | 0.0035 ** |
| sAHP delay after spike peak (ms) | $9.63 \pm 3.32$<br>(2.25–16.95) | $22.58 \pm 13.54$<br>(2.25–65.5) | $4.5 \times 10^{-12}$ *** |
| On cell mode spontaneous firing frequency<br>(Hz) | $20.79 \pm 18.74$<br>(0–90.53) | $5.47 \pm 9.22$<br>(0–35.68) | $1.2 \times 10^{-08}$ *** |
| Whole cell mode spontaneous firing<br>frequency (Hz) | $19.02 \pm 13.74$<br>(0–61.74) | $9.41 \pm 9.44$<br>(0–31.85) | $4.0 \times 10^{-06}$ *** |

GP<sub>STN</sub>, GP neurons projecting to the subthalamic nucleus; GP<sub>CPu</sub>, GP neurons projecting to the striatum; fAHP, fast afterhyperpolarization; sAHP, slow afterhyperpolarization; \*\*, $p < 0.01$ ; \*\*\*, $p$ $< 0.001$ . Statistical significance was examined using the Wilcoxon rank sum test. The range of each parameter is shown in parentheses.

Not all GP neurons exhibited inward photocurrent. 67/159 and 151/248 neurons did so for M1 and M2 stimulation, respectively (Fig. 2D). The locations of the GP neurons in which inward currents were observed were plotted (Fig. 2E). Consistent with the distribution of cortical axons, these locations were frequently around the center of the GP in coronal slices. Responsive neurons were similarly concentrated around the center of the GP along the rostro-caudal axis. Neurons responding to M1 terminal stimulation tended to be located in the dorsal GP, whereas those responding to M2 terminal stimulation were clustered in the ventral GP (Fig. 2E).

It is possible that the observed EPSCs were elicited by the STN via a di-synaptic circuit. However, we used coronal slices with an anteroposterior position of 0.6 mm rostral (r0.6) – 2.2 mm caudal (c2.2) to bregma, which did not include the STN (Paxinos & Watson, 2007). Bath application of the sodium channel blocker tetrodotoxin (TTX) at 1 µM completely prevented inward currents (Fig. 3A). Additional application of the potassium channel blocker 4-aminopyridine (4AP) at 1 mM recovered the currents to up to 60% of control on average (Fig. 3A), indicating that the current was monosynaptic (Gradinaru, Mogri, Thompson, Henderson, & Deisseroth, 2009; Shu, Yu, Yang, & McCormick, 2007). Moreover, a GABA_A_ receptor antagonist (gabazine, 20 µM) did not block the current (Fig. 3A); conversely, glutamate receptor antagonists (CNQX, 10 μM and AP-5, 20 μM) almost completely abolished the current (Fig. 3A). Thus, the inward current was mediated by glutamatergic excitatory postsynaptic currents (EPSCs). The latency (delay) of current onset after the photic stimulus in GP neurons did not differ from that in STN neurons (Fig. 3B1, 3B2). The 20–80% rise time and decay constant tended to be longer in GP neurons than in STN neurons (Fig. 3B3, 3B4). Taken together, these results indicated that the inward current observed in GP neurons was a monosynaptic, glutamatergic EPSC directly elicited by cortical terminals. Hereinafter, we refer to this current as an optically evoked EPSC (oEPSC).

**Fig. 3.**
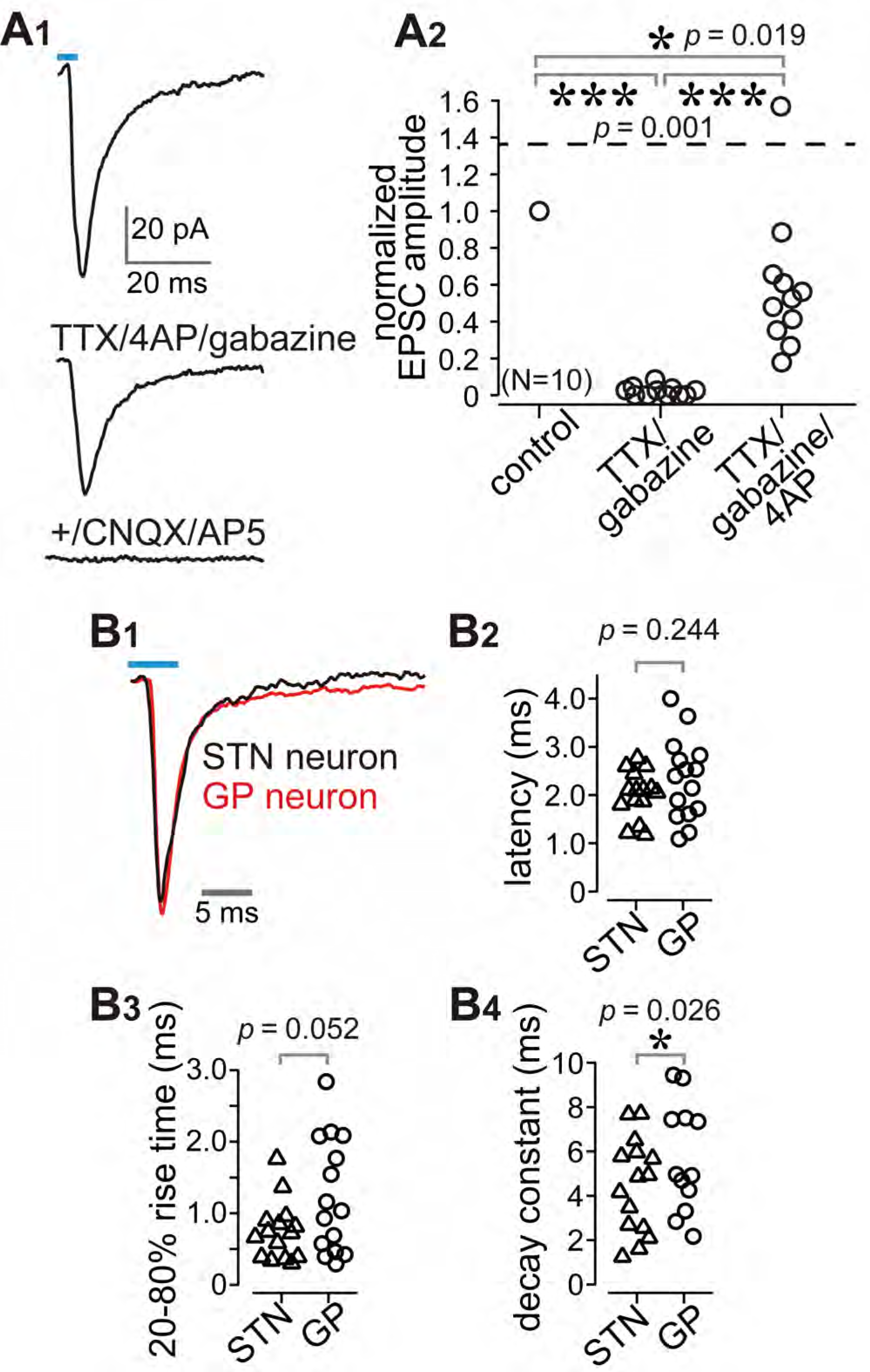
Cortico-pallidal connections are monosynaptic and glutamatergic. (A) The current induced by optogenetic stimulation of cortical terminals is monosynaptic and glutamatergic. (A1) Representative traces of pharmacological effects on the light-induced inward current. **Top**, no treatment; **Middle**, effects of TTX, 4AP, and gabazine; **Bottom**, effects of additional application of glutamate receptor antagonists (CNQX, AP5). (A2) A summary plot of pharmacological treatments (*N* = 10 GP neurons from 4 rats). (B) Comparison of the time courses of optogenetically evoked EPSCs (oEPSCs). (B1) Representative traces recorded from STN (black) and GP (red) neurons. The GP and STN neurons were not recorded simultaneously. (B2-4) Summary plots of oEPSC time courses in STN and GP neurons. (B2) The latency from light onset to oEPSC onset does not differ between STN and GP neurons (both *N* = 15 neurons from 6 rats). The rise time (B3) and the decay constant (B4) of oEPSCs (*N* = 15 GP neurons and 12 STN neurons). Statistical significance was examined by the Wilcoxon rank sum test. The STN and GP neurons were recorded from the different brain slices of the same animal.

### M1 and M2 preferentially innervate GP neurons projecting to the striatum than those projecting to STN

The results shown in Fig. 2C raised the question whether GP neurons have cell-type specific cortical innervation. To determine whether cortical innervation depends on the GP-neuron projection type, we conducted *in vitro* whole cell recordings from retrogradely labeled GP_STN_ or GP_CPu_ neurons. Occasionally, we recorded large GP neurons with no or very little spontaneous activity (*N* = 6, 1.20 ± 1.79 Hz during on-cell recording mode), which possessed a distinct action potential shape. Based on earlier reports (Bengtson & Osborne, 2000; Hernandez et al., 2015), these were most likely cholinergic neurons and were excluded from subsequent analysis, although they were found to be innervated by the cortex (*N* = 5/6). In addition to the molecular profiles, the electrophysiological properties of GP_CPu_ and GP_STN_ neurons also differed. GP_STN_ neurons usually showed spontaneous repetitive firing (∼20 Hz; Table 1), whereas many GP_CPu_ neurons were silent. Firing frequencies induced by depolarizing current pulses were higher in GP_STN_ than in GP_CPu_ neurons, and spike width was narrower in GP_STN_ neurons (Fig. 4B; see Table 1 for other electrophysiological parameters and quantitative comparisons). We discovered that GP_CPu_ neurons were more frequently innervated by motor cortex (51/62) than were GP_STN_ neurons (53/126). The oEPSC amplitude was also larger in GP_CPu_ neurons (Fig. 4D, E, F). The distribution of oEPSC amplitudes recorded from GP_CPu_ neurons seemed bimodal; the smaller-amplitude group was similar to GP_STN_ neurons (Fig. 4D). For comparison, MSNs (*N* = 11) and STN neurons (*N* = 18) were recorded. One-way ANOVA followed by post-hoc Tukey test revealed that the oEPSC amplitude was significantly larger in MSNs than in STN or GP neurons (for all combinations of comparisons, *p* < 9×10^−7^). The oEPSC amplitude of STN neurons was not significantly different from that of GP_STN_ (*p* = 0.109) or GP_CPu_ neurons (*p* = 0.999). GP_CPu_ neurons exhibited significantly greater amplitudes than did GP_STN_ neurons (*p* = 0.0021; Fig. 4D). Despite the larger EPSCs, we observed relatively hyperpolarized membrane potentials and lower membrane input resistances in MSNs compared with GP or STN neurons. The mean membrane potential (*V*_mean_) = −73.45 ± 5.3 mV and input resistance (*R*_in_) = 79.17 ± 28.38 MΩ for 10 MSNs; *V*_mean_ = −45.43 ± 7.47 mV and *R*_in_ = 253.14 ± 156.31 MΩ for 15 STN neurons; see Table 1 for GP neurons). Thus, the ease of induction of action potentials was lower in MSNs than in GP or STN neurons, at least in slice preparation. Actually, the median of rheobase current was 695 pA for MSNs (*N* = 10; range. 450–1415 pA), whereas 55 pA for STN neurons (*N* = 30; range, 10–545 pA) and 30 pA for GP neurons (*N* = 100; range, 5–250 pA).

**Fig. 4.**
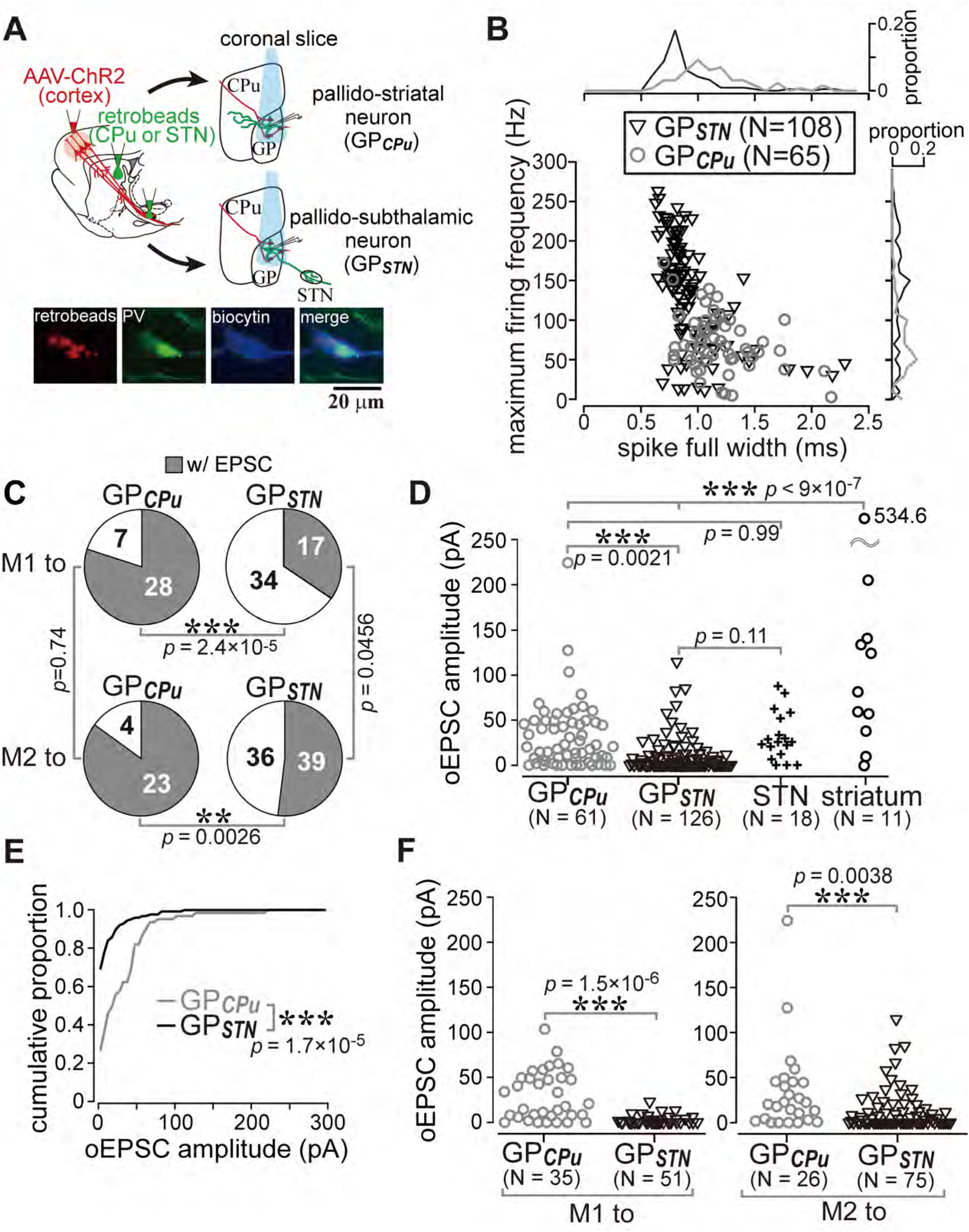
Cell type-dependent cortical innervation of pallidal neurons (A) Schematic of *ex vivo* recordings from retrogradely labeled GP neurons to investigate the effect of GP neuron projection type on cortical innervation. (B) Electrophysiological differences between GP neurons projecting to the striatum (GP_CPu_) and STN (GP_STN_) (see also Table 1). (C) The proportion of GP neurons innervated by M1 or M2 is correlated with projection type. GP_CPu_ neurons were more often innervated by either M1 or M2 than were GP_STN_. Significance was examined using Fisher’s exact test with a Bonferroni correction for multiple comparisons (***, *p* < 0.00025; **, *p* < 0.0025; *, *p* < 0.0125).(D) Amplitudes of oEPSCs in GP, STN, and striatal neurons. The amplitude in GP_CPu_ neurons is similar to that in STN neurons but smaller than that in striatal medium spiny neurons (MSNs). From each rat, CPu, GP and STN neurons were recorded in the same experimental session. Data obtained from M1 and M2 stimulation are summed. (E) Cumulative histograms of oEPSC amplitude in GP_CPu_ and GP_STN_ neurons. GP_CPu_ neurons exhibit a greater optically evoked EPSC (oEPSC) amplitude. (F) **Left**, amplitudes of oEPSCs induced in GP neurons by M1 terminal stimulation. The GP_CPu_ group shows larger oEPSC amplitudes than the GP_STN_ group. **Right**, amplitudes of oEPSCs induced by M2 terminal stimulation. Significantly larger oEPSCs were again recorded in the GP_CPu_ group (*p* = 0.034), but the difference is small. The neurons represented in Fig. 4 are not the same population as shown in Fig. 2, except for in Fig. 2C.

GP_CPu_ neurons were frequently innervated by either M1 (28/35) or M2 (23/27). In contrast, only a small fraction of GP_STN_ neurons received M1 (17/51) or M2 (36/75) inputs (Fig. 4C). The amplitude of M1-induced oEPSCs in the GP_CPu_ neurons was significantly larger than in the GP_STN_ neurons (*p* = 1.5×10^−6^ using the Wilcoxon rank sum test). This was also the case for M2-induced oEPSCs (*p* = 0.0038). Therefore, both motor areas preferentially innervated GP_CPu_ neurons, although GP_STN_ was more effectively innervated by M2 (Fig. 4C, 4F). Indeed, the mean oEPSC amplitude in GP_STN_ was larger with M2 stimulation than with M1 stimulation (*p* = 0.0028), but no significant difference between cortical sites was observed in GP_CPu_ neurons (*p* = 0.9595). M2 stimulation frequently evoked oEPSCs with initially smaller amplitudes, especially for the first light pulse, although repetitive light pulses at 10 Hz often augmented the oEPSC amplitude, similar to M1 stimulation. The paired-pulse ratio for second-to-first oEPSC in GP_STN_ neurons was 1.27 ± 0.81 for M2 stimulation (*N* = 38) and 1.45 ± 1.20 for M1 stimulation (*N* = 14); in the GP_CPu_ neurons, it was 1.25 ± 0.62 for M2 stimulation and 1.31 ± 0.79 for M1 stimulation.

To further identify characteristics of GP neuron types, using retrograde labeling of GP neurons and immunofluorescence against parvalbumin (PV), LIM homeobox 6 (Lhx6), and forkhead box protein 2 (FoxP2), we examined molecular profiles of GP neuron projection types in Wistar rats (Fujiyama, Nakano, et al., 2015). We confirmed that GP_CPu_ neurons frequently expressed FoxP2 or Lhx6, but not PV (*N* = 659 GP_CPu_ neurons in three sections from three rats; Fig. 5), in agreement with previous studies in mice (Dodson et al., 2015; Hernandez et al., 2015; Mastro et al., 2014; Mizutani, Takahashi, Okamoto, Karube, & Fujiyama, 2017), Long-Evans rats (Oh et al., 2017), and Sprague-Dawley rats (Abdi et al., 2015; H. Kita & Kita, 2001). The expression of FoxP2 and Lhx6 was almost mutually exclusive. Most PV(+) GP_CPu_ neurons co-expressed Lhx6. GP_STN_ neurons lacked expression of FoxP2 but expressed PV and/or Lhx6 (*N* = 727 GP_STN_ neurons in three sections from three rats; Fig. 5). Triple immunofluorescence combined with a single retrograde tracer injection into the striatum was conducted to further elucidate the molecular identity of GP_CPu_ neurons. The expression of Lhx6 (333/665) or FoxP2 (328/665) was again almost mutually exclusive. Only a small fraction (14.7%) of Lhx6-expressing neurons co-expressed PV (49/333) (Fig. 5B). Occasionally, double-retrogradely labeled neurons, namely bi-directional projecting GP neurons (GP_Bi_), were observed (*N* = 113). Lhx6 was expressed in most GP_Bi_ neurons (61/69), PV less frequently (19/72), and FoxP2 rarely (2/85). GP_Bi_ neurons that expressed PV co-expressed Lhx6 in most cases examined (5/6). Therefore, GP_STN_ neurons comprised PV(+) and/or Lhx6(+) prototypic neurons, whereas GP_CPu_ neurons comprised arkypallidal neurons expressing FoxP2 and prototypic neurons expressing Lhx6.(Fujiyama, Nakano, et al., 2015; Mallet et al., 2012). Importantly, these data insist that selective electrophysiological recording from Lhx6(+) neurons can be accomplished by targeting GP_Bi_ neurons. As a result, we found that GP_Bi_ also received cortical inputs (7/10 for M1 and 7/9 for M2; *N =* 3 rats for each), and most of them exhibited an oEPSCs with small amplitude (Fig. 5C). Actually, the frequency distributions of oEPSC amplitudes significantly differ between GP_Bi_ and GP_CPu_ neurons (Fig. 5D; *p* = 2.2×10^−16^ by the Kolmogorov-Smirnov test). It suggests that arkypallidal neurons, which do not send axons to the STN, could be a principle target of cortico-pallidal innervation.

**Fig. 5.**
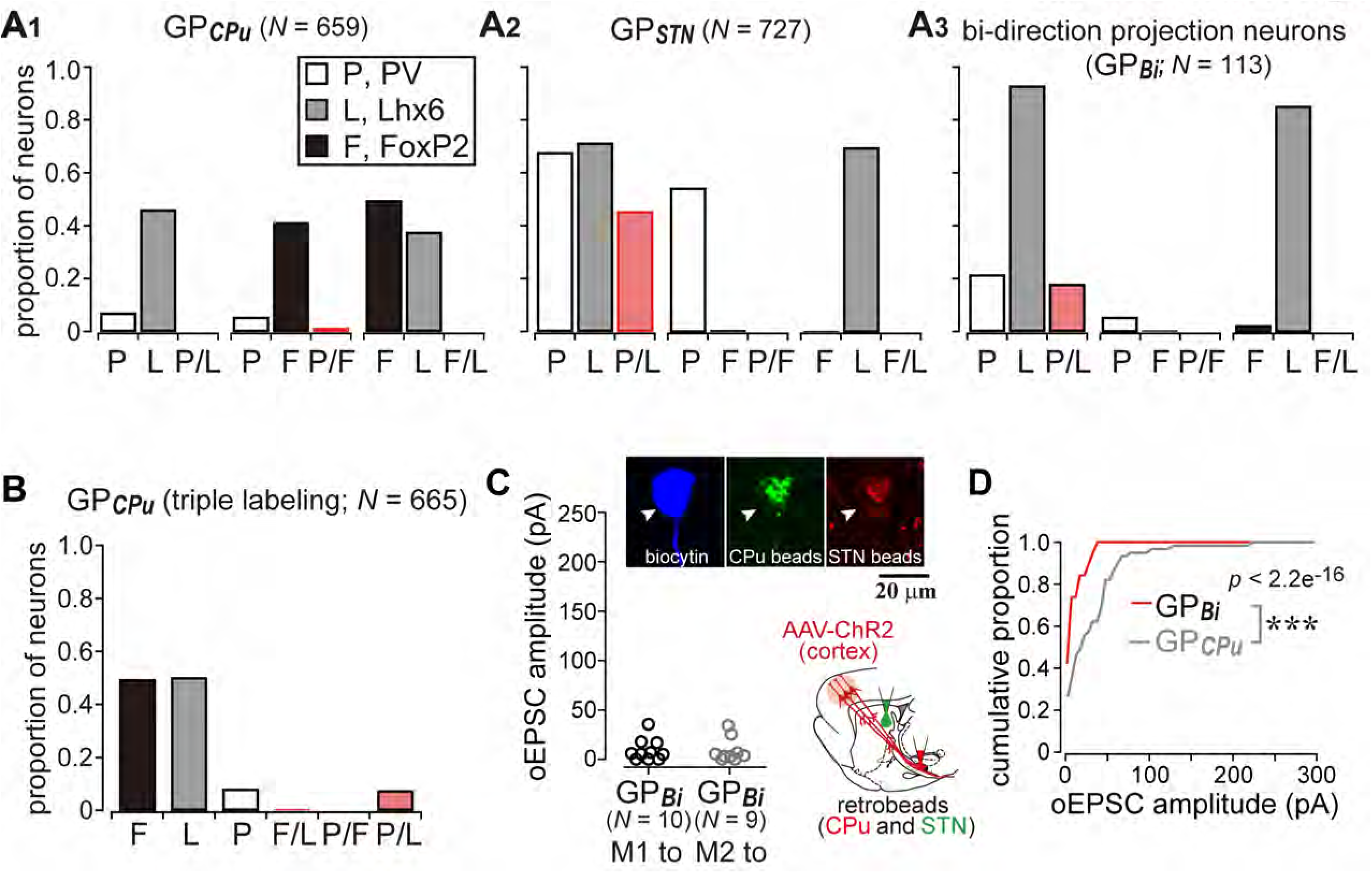
Cortical inputs on bi-directional projection GP neurons (GP_Bi_) Two distinct retrograde tracers were injected into the STN and striatum (CPu), respectively. (A) Three combinations of double immunofluorescence (PV/Lhx6, PV/FoxP2, and Lhx6/FoxP2) were applied (*N* = 3. rats). CPu-projecting GP neurons (GP_CPu_) expressed either FoxP2 or Lhx6 exclusively (A1), whereas STN-projecting GP neurons (GP_STN_) were mainly composed of PV- and/or Lhx6- expressing neurons (A2). (A3) GP neurons projecting to both the CPu and STN (GP_Bi_) frequently expressed Lhx6, but not FoxP2. (B) Triple immunofluorescence for GP_CPu_ neurons (*N* = 2. rats). Only a single retrograde tracer was injected into the striatum. (C) oEPSC amplitude of GP_Bi_ neurons (*N* = 3. rats for each M1 and M2 labeling). Most of GP_Bi_ exhibited small amplitude of oEPSCs. Green and red retrobeads were injected into the CPu and STN, respectively. Confocal images of a biocytin filled GP_Bi_ neuron (arrowheads) are shown. AAV-labeled cortical axons also show red fluorescence. (D) Cumulative histograms of oEPSC amplitude in GP_Bi_ and GP_CPu_ neurons. The distributions are significantly different (*p* = 2.2×10^−16^ by the Kolmogorov-Smirnov test). The histogram of GP_CPu_ neurons is the same as shown in Fig. 4E.

### M1 and M2 differentially innervate striatal subregions

The preceding results suggest that the two cortical motor areas project differentially to the GP. In rodents, cortical pyramidal cells send axon collaterals to multiple nuclei of the BG which, in turn, are themselves interconnected. Thus, M1 and M2 projections to nuclei other than the GP should involve information processing in cortico-basal ganglia circuitry (Lerner et al., 2015). To quantify the cortical axon distribution onto striatal molecular subregions, we employed AAV vector injection into the motor area followed by immunostaining for either CB, or μ-opioid receptor (MOR) which is a marker of the striosomes (Crittenden & Graybiel, 2011; Kincaid & Wilson, 1996; J. B. Smith et al., 2016). CB was preferentially expressed in the medial and ventral portion of the striatum and faintly in the dorsolateral striatum (Fig. 1-figure supplement 1) (H. Kita & Kita, 2001; Wouterlood, Hartig, Groenewegen, & Voorn, 2012). M1 axons were mostly confined to CB(-) subregions of the striatum (Fig. 6A), whereas M2 axons innervated both CB(-) and CB(+) striatal subregions (Fig. 6B). We quantified the axon distribution in CB(+) and CB(-) striatal subregions for both M1 and M2 axons (*N* = 3 rats for each subregion; 12 sections for M1 and 13 sections for M2). The proportion of CB(+) pixels that contained axons was calculated in the striatum. M2 axons were found in 37.2 ± 9.0% of CB(+) pixels, and M1 axons were found in only 15.3 ± 7.6% of CB(+) pixels (*p* = 0.000012 by Wilcoxon rank sum test; Fig. 6C1). To evaluate the relative strengths of the innervation, the brightness of the fluorescence in each pixel in the regions of interest (ROIs) were measured. The median pixel intensity in CB(-) ROIs was normalized to that of CB(+) ROIs. As shown in Fig. 6C2, the normalized fluorescence intensity in CB(-) striatum was ∼1.6-times higher for both M1 (1.61 ± 0.39) and M2 (1.56 ± 0.28) axons. These results indicated that M2 axons innervated CB(+) striatum more frequently than did M1 axons, although CB(-) ROIs were more densely innervated by both M1 and M2 than were CB(+) ROIs.

**Fig. 6.**
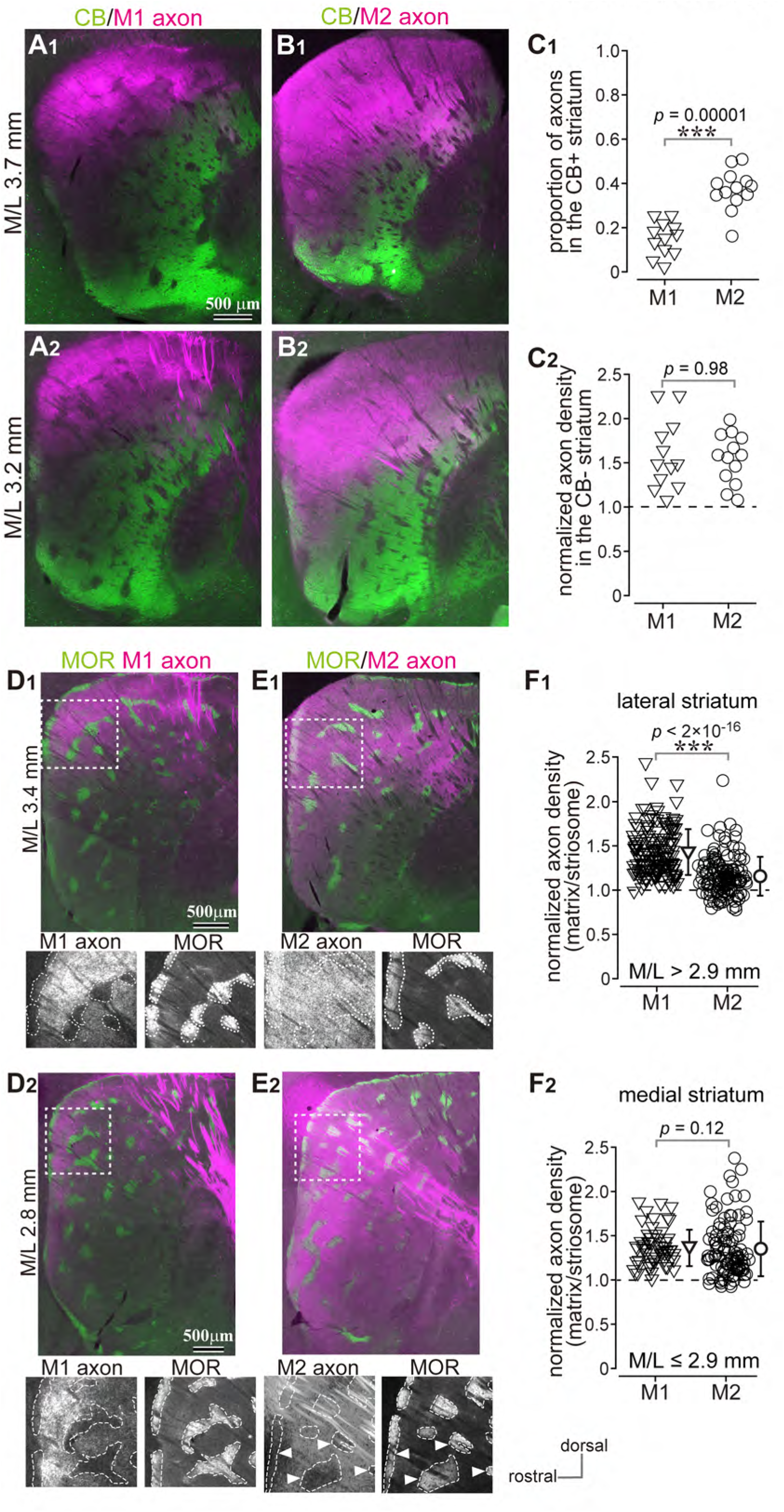
Distribution of cortico-striatal axons in striatal subregions (A) Distribution of M1 cortico-striatal axons (magenta) in relation to calbindin (CB) expression (green). Sagittal sections at two mediolateral (M/L) coordinates are shown (A1, M/L 3.7 mm; A2, M/L 3.2 mm). Note that M1 axons were densely distributed in the dorsolateral striatum, where CB expression was weak. (B) M2 corticostriatal axons. M2 axons were distributed in the CB(+) and dorsal CB(-) subregions at M/L 3.7 mm (B1) and M/L 3.2 mm (B2). (C) Quantitative comparison of axon distributions between M1 and M2 (for each *N* = 3 rats). (C1) Proportion of axon-containing pixels in the CB(+) striatum. M2 axons more frequently innervated the CB(+) striatum than did M1 axons. (C2) Normalized fluorescence intensity in the CB(-) striatum, which reflects axon density, was not significantly different between M1 and M2. The dotted line at 1.0 indicates axon density in the CB(+) striatum. (D) Distribution of M1 axons in the striosome/matrix subregions of the striatum. Upper panel, M1 axons (magenta) are overlaid with μ-opioid receptor (MOR) immunofluorescence (green). Dotted area in the panel is magnified in the bottom two panels. The fluorescence of M1 axons (**bottom left**) and MOR (**bottom right**) is shown in grayscale. MOR immunopositive striosomes are contoured by white dotted lines. Note that M1 axons seemingly avoided the striosomes in both lateral (D1, M/L 3.4 mm) and medial (D2, M/L 2.8 mm) sections. (E) Distribution of M2 axons in the striatum, as described in (D). Note that in lateral sections (M/L 3.4 mm, E1), M2 axons were uniformly distributed in both matrix and striosome, whereas in medial sections (M/L 2.8 mm, E2), M2 axons were observed in dorsal but not ventral striosomes (arrowheads). (F) Normalized fluorescence intensity in the matrix (*N* = 3 rats for each M1 and M2 labeling). M1 axons were more densely distributed in the matrix in lateral sections (M/L > 2.9 mm, F1), whereas both M1 and M2 preferentially innervated the matrix in medial sections (M/L ≤ 2.9 mm, F2). The dotted black line represents axon density in the striosomes. The sections from the same rat were used for either CB or MOR immunostaining.

Comparing the data in Fig. 6A and 6B shows that M1 and M2 axons exhibited differing innervation of the striosome. Based on MOR immunofluorescence, M1 axons appeared to preferentially innervate the matrix, and axon labeling in the striosomes was faint (Fig. 6D). In contrast, M2 axons were almost equally distributed in both the striosome and matrix (Fig. 6E), as previously reported for mouse (J. B. Smith et al., 2016). We also observed topographical differences in striosome preference. In medial sections (M/L ≤ 2.9 mm), the striosomes in the ventral part of the dorsal striatum were not densely innervated by M2 axons (arrowheads in Fig. 6E2; see also Fig. 1-figure supplement 1). We quantified relative preference by normalizing fluorescence intensity in the matrix to that of the striosomes (Fig. 6F). For M2, the matrix preference was only 1.17 ± 0.22 (*N* = 3 rats, 126 ROIs) in lateral sections (M/L > 2.9 mm). In contrast, the matrix preference for M1 was 1.45 ± 0.26 (*N* = 3 rats, 155 ROIs), a significantly greater preference for the matrix than that of M2 axons (*p* < 2.2×10^−16^ by Wilcoxon rank sum test; Fig. 6F1). In medial sections (M/L ≤ 2.9 mm), both M1 and M2 preferred the matrix similarly, i.e., 1.37 ± 0.20 for M1 (*N* = 60 ROIs) and 1.36 ± 0.31 for M2 (*N* = 106 ROIs; Fig. 6F1). Taken together, these results show that striosomes in the dorsal striatum were selectively innervated by M2, especially in the lateral striatum.

### Cortico-subthalamic (STN) projections from frontal cortical areas

Cortico-STN projections are topographically organized depending on cortical area of origin, especially in primates (Haynes & Haber, 2013; Nambu, Takada, Inase, & Tokuno, 1996; Nambu, Tokuno, Inase, & Takada, 1997). Retrograde tracer injection into the STN labeled L5 cortical neurons in a wide range of areas, among them, many neurons were found in M1, M2, and the lateral orbitofrontal area (LO) in the frontal cortex (Fig. 7-figure supplement 1). We also observed that M1 and M2 had different topographies of cortico-STN projection. M1 axons were concentrated in the central part of the STN along the anteroposterior and dorsoventral axis (Fig. 7A, B). The density of M1 axons was low around the rostral and caudal STN. The axon distribution in STN was similar between rostral and caudal M1 areas (Fig. 7A, B). Conversely, M2 axons were densely distributed in the dorso-anterior and ventro-posterior portions of the STN (Fig. 7C). LO provided dense axonal input to the rostral part of the STN (Fig. 7D). We quantified the preferential axon distribution in STN along the anteroposterior (Fig. 7E1) and dorsoventral axes (Fig. 7E2). Significant differences were observed among M1, M2, and LO (*p* < 0.05 by the Kruskal-Wallis test followed by multiple comparisons based on the Fisher Exact test with a Bonferroni correction for multiple comparisons). Of note, the topographic distribution was not clearly segregated; rather, a substantial overlap of axons from different frontal cortical areas was observed (Fig. 7F).

**Fig. 7.**
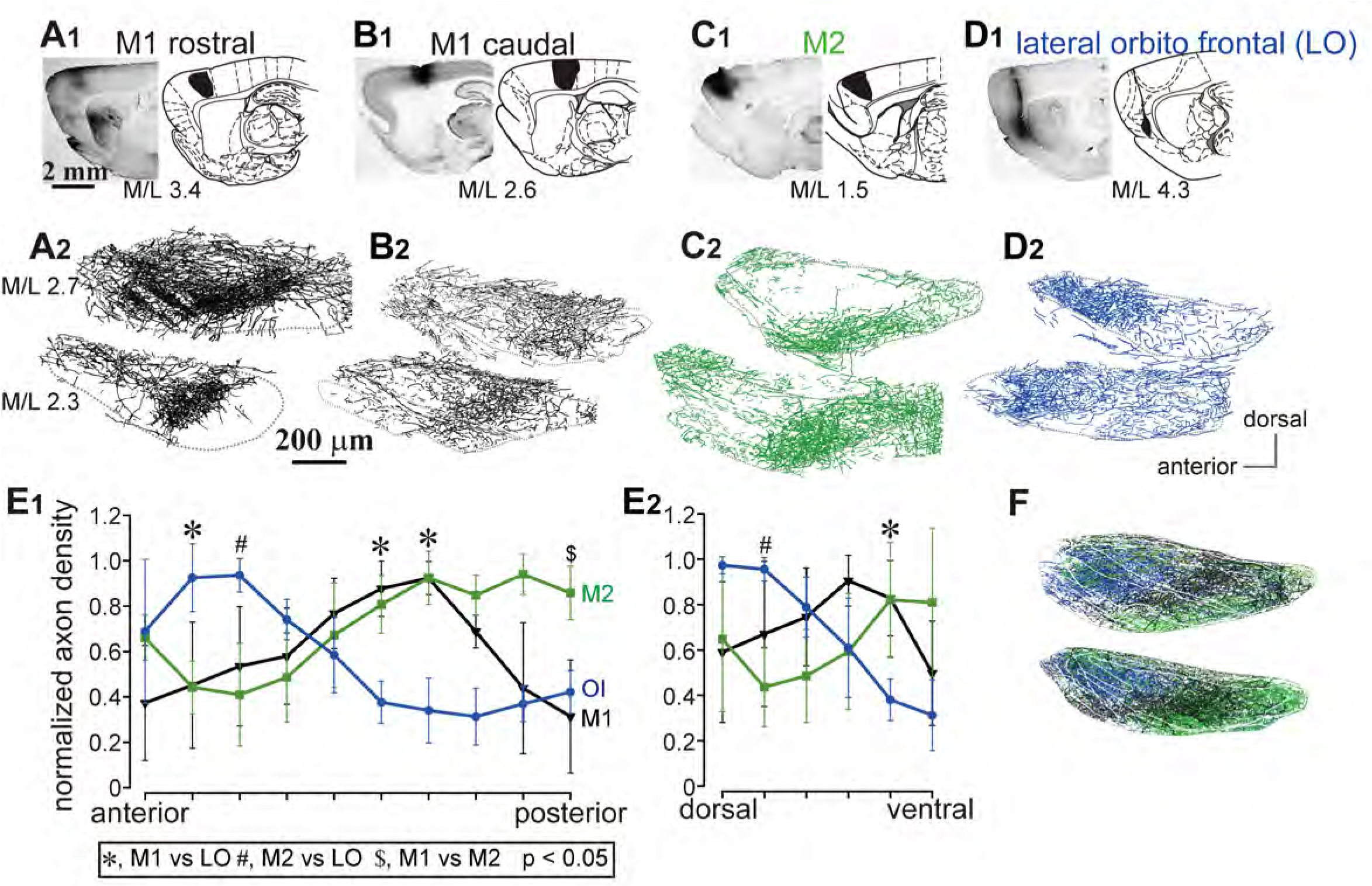
Distribution of cortico-subthalamic axons Images and drawings of BDA injection sites into rostral M1 (A1), caudal M1 (B1), M2 (C1), and lateral orbitofrontal area (LO). LO injections often extended to the insular area. However, additional experiments revealed that the insular cortex did not send many axons to the STN, as previously reported (Tsumori et al., 2006). Cortical axons in STN are represented in two sagittal planes (M/L 2.3 and 2.7 mm) for M1 (A2, B2), M2 (C2), and LO (D2). The gray dotted line indicates the boundary of the STN. (E) Normalized distribution of cortical axons in the STN along the A/P (E1) or D/V axis (E2). Data from sections M/L 2.3 mm and 2.7 mm are pooled (*N* = 3 rats for M1, *N* = 2 for M2, and *N* = 2 for LO). (F) Merged traces of axons from M1, M2, and LO (*N* = 2 rats for each cortex). M1 (black) terminated in the central part of the STN, whereas M2 axons (green) were dense in the posterior-ventral portion.

### Neural projections among the striatum, GP, and STN

L5 PT-type neurons, candidates for the origin of the cortico-pallidal innervation, are known to project to multiple brain nuclei including the striatum and STN (T. Kita & Kita, 2012; Shepherd, 2013; Shibata et al., 2018) This suggests that when the GP receives cortical excitation, the striatum and STN are co-excited by the same source. Therefore, to elucidate the functional relevance of cortico-pallidal pathways, it is important to examine the relationship between their axon distribution and that of other, related intra-basal ganglia projections to elucidate functional relevance of cortico-pallidal pathways. For bi-directional projections between GP and the striatum, CB expressions in the GP and striatum are correlated, such that the CB(+) striatum (medio-ventral striatum) projects to the outer edges of the GP, where CB immunoreactivity is obvious due to innervation by CB(+) striatal axon terminals (H. Kita & Kita, 2001). In contrast, the dorsolateral striatum which has sparse CB expression, innervates the central part of the GP, which lacks CB immunoreactivity (H. Kita & Kita, 2001; Rajakumar, Elisevich, & Flumerfelt, 1994; Rajakumar, Rushlow, Naus, Elisevich, & Flumerfelt, 1994). We confirmed these findings using tracer injections into the striatum (Fujiyama et al., 2011; Kawaguchi, Wilson, & Emson, 1990; Wu, Richard, & Parent, 2000) (Fig. 8). Thus, the central GP can receive direct cortical excitation (Fig. 1, 2; Fig. 1-figure supplement 1) and iMSN innervation from the dorsolateral striatum, where motor cortex preferentially projected (Fig. 6). It suggests convergence of motor cortical information. Projections from the GP to the striatum are likely to obey the same rule (Fig. 8). For GP-to-STN projections, we found topographic projection patterns as shown in Fig. 9. The rostral GP, expressing CB, projected to the rostral part of STN, whereas the central GP, lacking CB expression, projected to the central part of STN. Because the rostral STN received M2 projection and the central STN received M1 projection, again, motor cortical information can converge in the STN subregions.

**Fig. 8.**
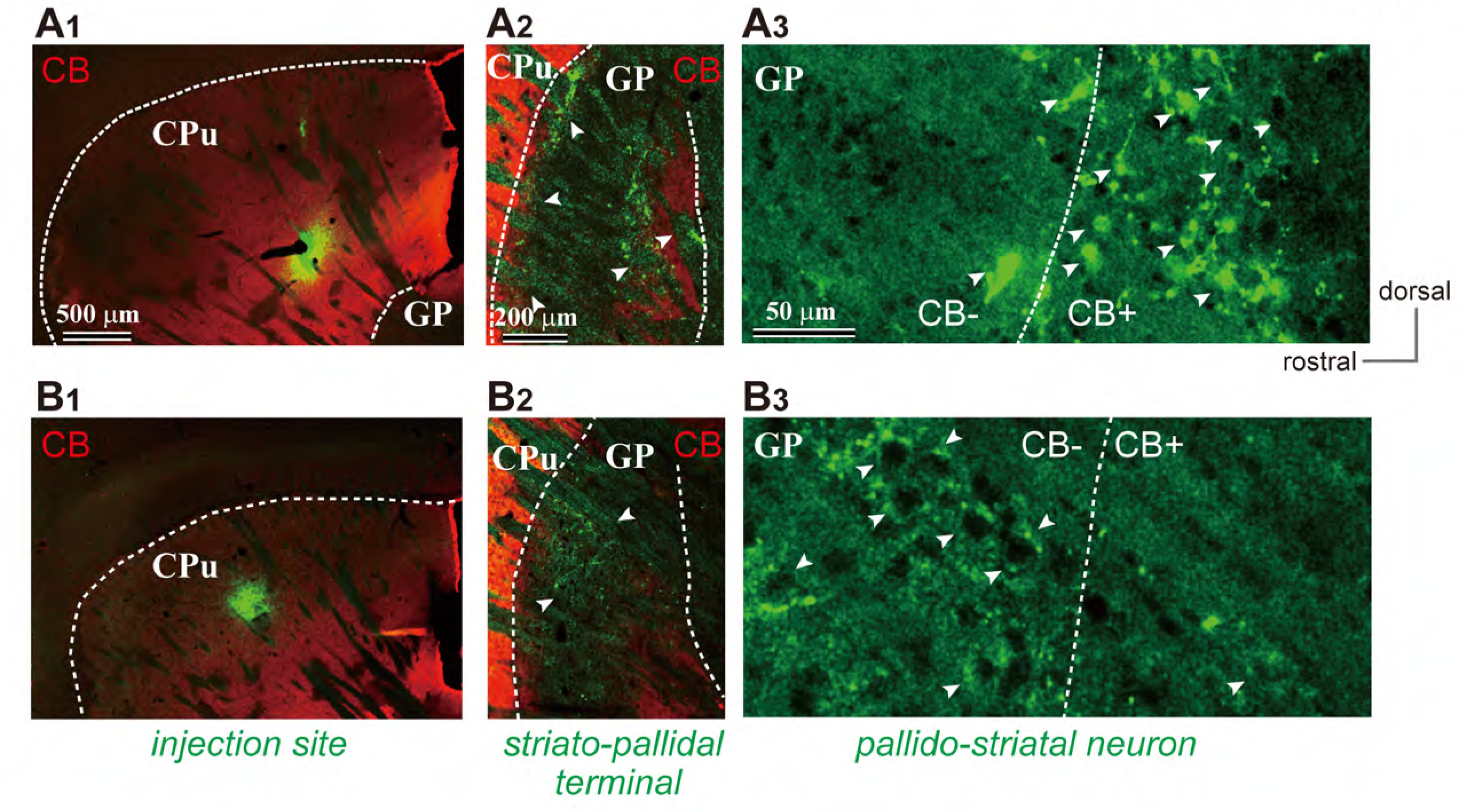
Reciprocal projections between the striatum and GP Cholera toxin subunit B conjugated with Alexa-Fluor-488 (CTB488) was injected into the striatum (CPu), either the CB(+) (A1) or CB(-) (B1) subregions. CTB488 can be transported in both anterograde and retrograde directions. In the GP, anterogradely labeled terminals of medium spiny neurons (MSNs) (arrowheads in A2 and B2) and labeled GP neurons projecting to the CPu (arrowheads in A3 and B3) were observed. Reciprocal projections between CPu and GP correlated with CB expression. Dotted lines show the boundary of the striatum (A1, B1), the GP (B2, C2), and CB(+)/CB(-) subregions of the GP (A3, B3). Representative images from one rat for each are shown.

**Fig. 9.**
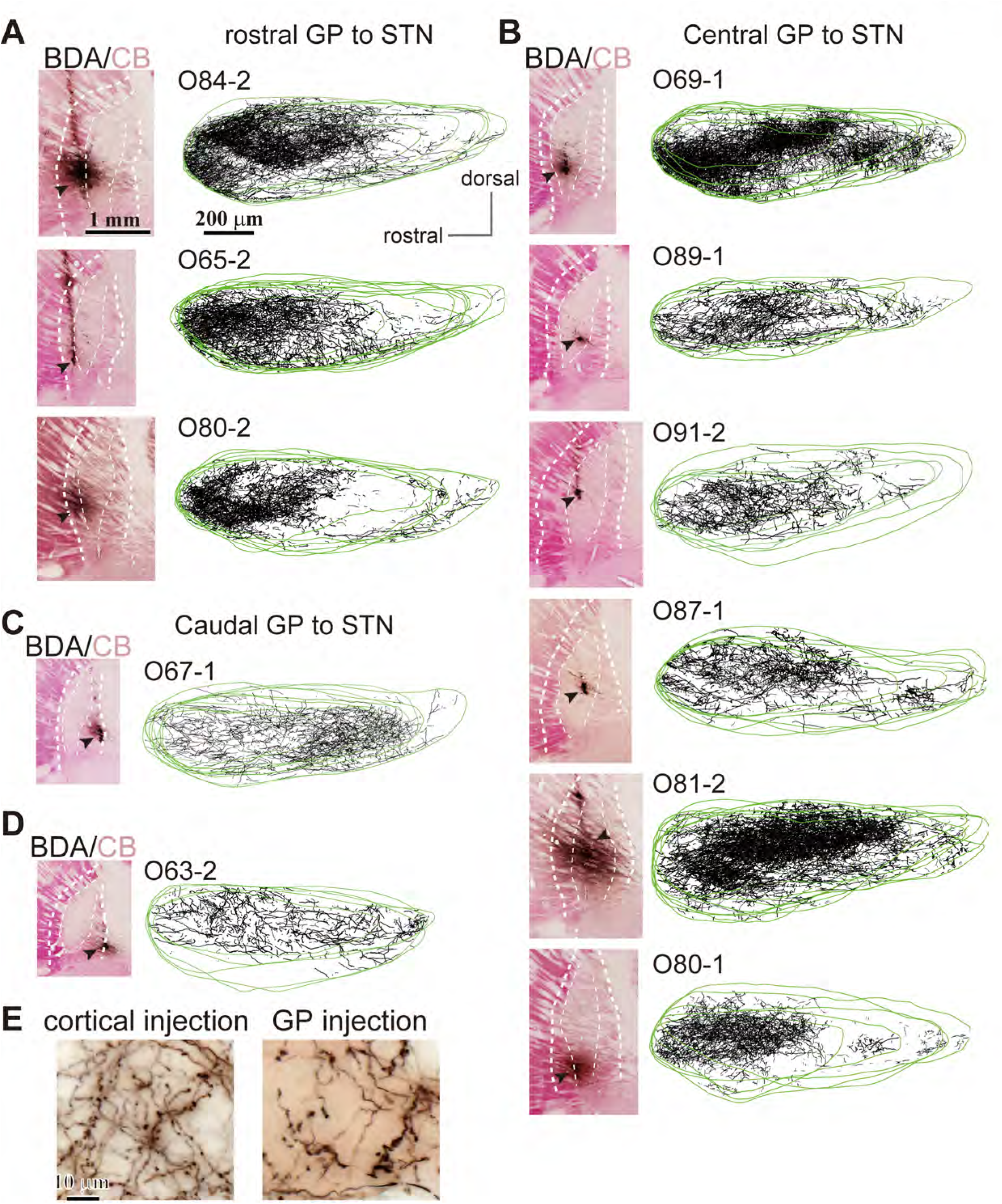
Projections from GP to STN (A), (B), (C) **Left**, Images of BDA injection sites in the GP subregions. Sections containing the injection core (arrowheads) were counterstained for CB (magenta). Thick dotted lines indicate the borders of the GP, and thin dotted lines indicate the borders of CB(+) and CB(-) subregions of the GP. **Right**, Drawings of labeled GP axons in the STN. Labeled axons (black) in the STN were traced over STN contours (green). Rostral GP projected to rostral STN (A; *N* = 3 rats) and central GP to central STN (B; *N* = 6 rats). (C) In one case, we observed caudal GP projections to caudal STN (*N* = 1 rat). (D) Injection around the border between the GP and the internal capsule (*N* = 1 rat). Axons in the STN are uniformly distributed in the STN, probably due to labeling of fibers of passage from the entire GP. Four to eight STN sections are overlaid for each case. Scales shown in A pertain also to B and C. The labeled axons do not include striatal MSN axons because the striatum does not project to the STN. Passing fibers of cortico-STN projections may also be labeled with tracer injections into the GP; however, this is unlikely in our experiments because the morphology of axonal boutons in the STN differed between cortical and GP injections (E), as reported in monkeys (Shink & Smith, 1995).

## Discussion

To summarize, we here report that direct motor cortical innervation of GP neurons is as dense as that of STN neurons. Pallidostriatal neurons were more frequently and heavily innervated by the cortex than were pallidosubthalamic neurons. We also investigated the distribution of cortical projections onto basal ganglia subregions, which likely affects the functions of the cortico-pallidal pathway, as discussed subsequently (see Fig. 10 for the schematic). To date, fast excitation observed in GP following cortical stimulation has been considered disynaptic excitation via the STN (Nambu et al., 2000). This does not fully contradict the present findings, since a larger population of GP neurons, GP_STN_ (or prototypic neurons), was not the main target of the motor cortex. In addition, it is possible that traditional extracellular unit recordings are biased towards neurons with relatively higher firing frequency, which include GP_STN_ neurons (Table 1). A question that remains is whether cortico-pallidal projections exist in primates including humans, as suggested (Milardi et al., 2015; Y. Smith & Wichmann, 2015).

**Fig. 10.**
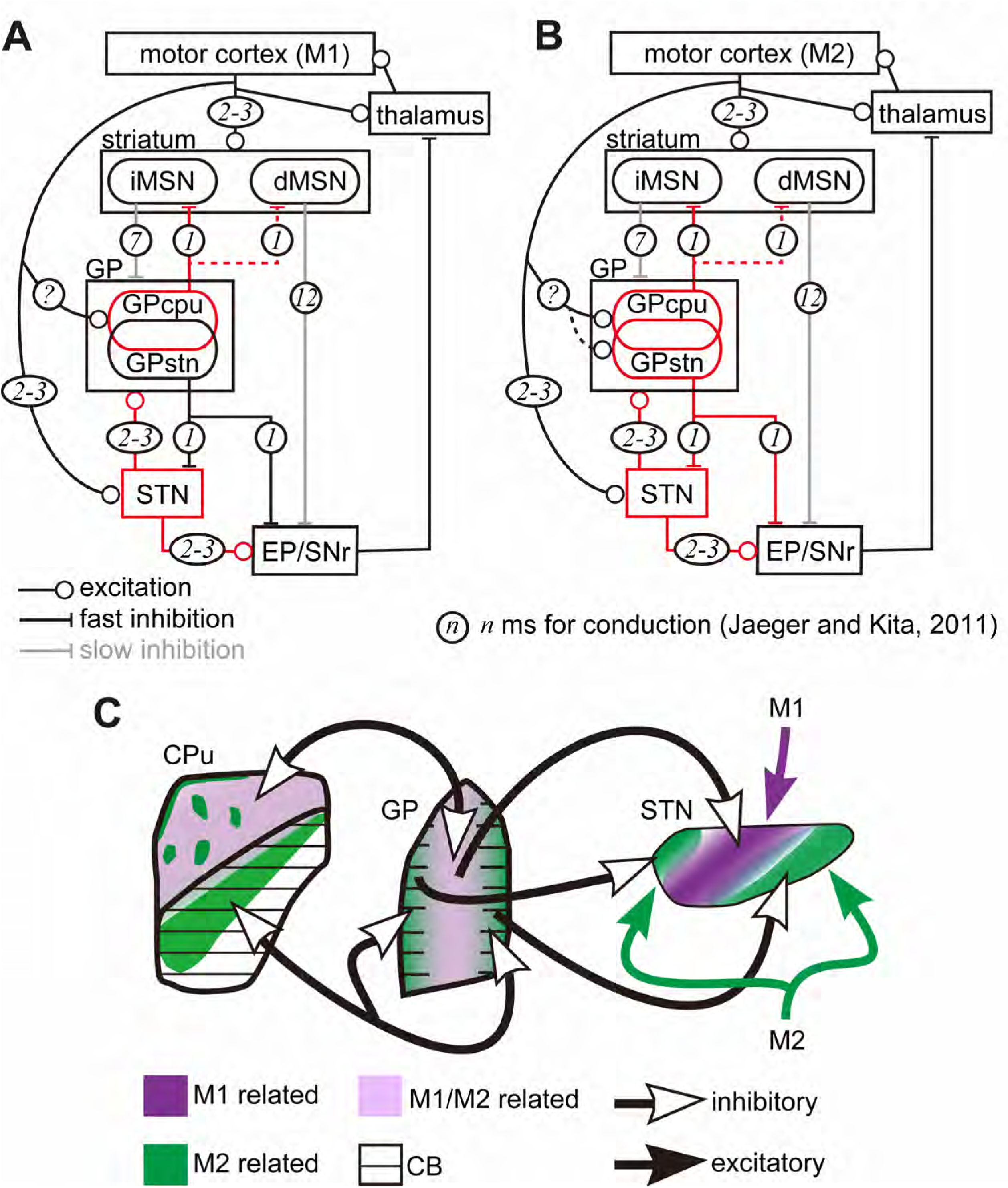
Schematic drawings of the cortex-basal ganglia circuitry Diagram of cortico-basal ganglia-thalamus circuitry. The dotted lines represent the relatively weak innervation reported in this study for cortex to GP_STN_, and for GP_CPu_ to dMSNs (Glajch et al., 2016). The numbers in circles indicate estimated conduction times in millisecond (ms) (Jaeger & Kita, 2011). (A) M1 activation induces excitation in the striatum, STN, and GP_CPu_. Due to the electrophysiological properties of the neurons, STN and GP may be activated faster than striatal neurons. (B) M2 activation conveys additional excitation to the GP_STN_ neurons. Possible information flow related to the present study is shown by red lines. Note that inhibition from the GP is here considered faster (1 ms) than excitation from the STN (2–3 ms) (Jaeger & Kita, 2011). The actual timing of spike activity depends on neuron type, and excitation/inhibition interactions are complex. (C) Schematic of connections among the basal ganglia nuclei with relation to M1 and M2 innervation.

### Innervation of the basal ganglia by the frontal cortex

Topographic cortical projections are well established for the striatum and STN (Heilbronner, Meyer, Choi, & Haber, 2018; Hooks et al., 2018; Janssen et al., 2017; Mathai & Smith, 2011; Nambu, 2011; J. B. Smith et al., 2016). We also demonstrated topographic differentiation of cortico-striatal projections between M1 and M2 with distinct molecular-subdomain preferences (Fig. 6). For other frontal or associative areas, we observed that LO innervated the ventromedial part of the striatum with preference for the striosomes, whereas the cingulate cortex (Cg) projected to the dorsomedial striatum and avoided the striosomes, consistent with an earlier report (Friedman et al., 2015). For cortico-STN projections, M1 and M2 projected densely to the central-to-caudal portions of the STN, whereas LO and Cg projected to the rostromedial and caudomedial STN, respectively (Supplementary Fig. 1, 2). Cortico-pallidal projections were also topographically organized; M1 and M2 axons were densely distributed in the CB(-) central part of the GP (Fig. 1-figure supplement 1). Cg axons projected to the GP in a similar manner. In contrast, LO provided fewer pallidal collaterals (Fig. 1, Supplementary Fig. 1, 2). Thus, each frontal cortical area likely has its own regions of interest in each BG nucleus. In addition, CB(-) GP was interconnected with CB(-) striatum (Fig. 8; Fujiyama et al., 2011; H. Kita & Kita, 2001), and projected to the central part of the STN (Fig. 9), which received projections from motor cortices, especially M1. Similar topographic interconnections between STN and GP have been reported in primates (Shink, Bevan, Bolam, & Smith, 1996) and rodents (Baufreton et al., 2009; but see also Canteras, Shammah-Lagnado, Silva, & Ricardo, 1990; van Dijk et al., 2016). Moreover, thalamic inputs also differentiate between CB(+) and CB(-) GP (Y. Smith, Raju, Pare, & Sidibe, 2004), and GP neurons form topographic pallido-pallidal connections dependent on the location of the cell bodies (Sadek, Magill, & Bolam, 2007). Therefore, topographic interconnections among the cortex-striatum-GP-STN likely contribute to differential integration of neural signals.

### Possible effects of cortical innervation in GP

Our findings may explain the heterogeneous activity of GP neurons during movement (Arkadir, Morris, Vaadia, & Bergman, 2004; DeLong, 1971; Dodson et al., 2015; Goldberg & Bergman, 2011; Mink & Thach, 1991a, 1991b; Turner & Anderson, 1997). The firing phases of prototypic neurons (GP_STN_) and arkypallidal neurons corresponding to FoxP2(+) GP_CPu_ differ with regard to cortical activity (Abdi et al., 2015; Mallet et al., 2012; Mallet et al., 2016). This may be due to biased cortical innervation of GP_CPu_ (Fig. 4C, D, E, F). The axon terminal density of cortico-pallidal projections was sparser than cortico-striatal ones, but as dense as cortico-subthalamic ones. The neural connection in the basal ganglia is usually sparse; for example, even in the connection between STN and GP, which is clearly functional, only 1–2% of all GP neurons converge onto a single STN neuron and vice versa (Baufreton et al., 2009; Goldberg & Bergman, 2011). Moreover, even if excitatory synaptic inputs did not reach action potential threshold, the timing of subsequent action potentials is still altered (Ermentrout, 1996; Schultheiss, Edgerton, & Jaeger, 2010). Expression of sodium channels characteristic of GP neurons can boost excitatory synaptic inputs (Edgerton, Hanson, Gunay, & Jaeger, 2010; Hanson, Smith, & Jaeger, 2004). Thus, the relatively sparse cortical innervation to the GP_STN_ neurons could nevertheless affect a small but specific component of the basal ganglia circuitry.

Most neighboring GP neuron pairs do not exhibit real correlation (Bar-Gad, Heimer, Ritov, & Bergman, 2003; Goldberg & Bergman, 2011; Stanford, 2003), even for pause periods (Elias et al., 2007). Excitatory inputs may synchronize GP neurons more effectively, as GP neurons tend to align with STN inputs rather than with striatal inputs (Goldberg, Kats, & Jaeger, 2003). Cortico-pallidal inputs probably precede STN-GP inputs and thus strengthen STN-GP coupling (Bevan, Magill, Hallworth, Bolam, & Wilson, 2002; Bevan, Magill, Terman, Bolam, & Wilson, 2002; Parent & Hazrati, 1995b), thereby affecting the oscillatory phase and synchronization of GP neurons.

### Potential roles of the cortico-pallidal pathway

Mallet et al. (2012) reported that both MSNs and interneurons are innervated by arkypallidal neurons. Npas1(+) neurons, which overlap with Lhx6(+) and FoxP2(+) GP_CPu_ neurons, synapse onto iMSNs and dMSNs. The average inhibitory postsynaptic current amplitude is significantly larger in iMSNs (Glajch et al., 2016). In turn, iMSNs inhibit all GP neuron types equally (Hernandez et al., 2015). GP_STN_ is also activated by M2 terminals. A substantial difference among GP_STN_ cell types has been reported for innervation of the SNc (Mastro et al., 2014; Oh et al., 2017). Based on our results suggesting that M2 but not M1 project to the dorsolateral striosomes (Fig. 6) and the fact that striosomal dMSNs innervate the SNc (Fujiyama et al., 2011; Gerfen, 1985), M2-driven pathways may modulate dopamine signaling via the striatum and GP_STN_. Recently, Viana-Magno et al. (Viana Magno et al., 2019) revealed that M2 activation can relieve the motor dysfunction of Parkinson’s disease in mice. The neural circuitry proposed here is highly likely to contribute to such an effect.

As shown in Fig. 1-figure supplement 2, the cortico-pallidal pathway likely originates from PT neurons, which also issue axon collaterals to STN (Shepherd, 2013; Shibata et al., 2018). In addition, dense labeling of L5 neurons following injection of retrograde tracer in STN (Fig. 7-figure supplement 1) suggested that STN-projecting neurons are unlikely to form a specific population of PT neurons in rodents (Shibata et al., 2018). Thus, once M1 activates, cortical signals are transmitted to the striatum, STN, and GP (Fig. 10). Due to their membrane properties, STN and GP neurons will be activated more effectively and earlier than striatal neurons (Jaeger & Kita, 2011). Since GP_CPu_ neurons are activated by direct M1 inputs, they may exhibit more activity than GP_STN_ neurons and be more in phase with cortical excitation. In contrast, GP_STN_ neurons are not strongly activated by M1. The M1-pallidal pathway therefore is unlikely to counteract hyperdirect inputs onto STN neurons. Because the latency and strength of oEPSCs in GP_CPu_ neurons were almost equal to those in STN neurons (Fig. 6, 11), STN and GP neurons may be excited simultaneously. Inhibition of MSNs by GP_CPu_ could counteract corticostriatal excitation and weaken iMSN activity. Decreased iMSN activity may reduce inhibition of GP neurons, thereby augmenting the inhibition of STN, EP, and SNr. Operation of this trisynaptic circuit may require a long duration; thus, the M1 cortico-pallidal pathway may act as a delayed terminator of the hyperdirect pathway. GP_CPu_ can also inhibit dMSNs, which in turn disinhibit the EP/SNr. If iMSN activity is suppressed by GP_CPu_, what controls cortico-GP_CPu_ excitation? The most parsimonious explanation is that decreased cortico-pallidal activity can weaken GP excitation. In addition, GP_CPu_ inputs onto iMSNs will not completely block cortico-striatal excitation, but may delay it. It is noteworthy that striatal neurons receive contralateral cortical excitation as well as ipsilateral input (Wilson, 1986, 1987), whereas the GP and STN receive only ipsilateral excitation (Fig. 1-figure supplement 2). It is also possible that axon collaterals of dMSNs inhibit the GP (Fujiyama et al., 2011; Kawaguchi et al., 1990; Lévesque, Bédard, Cossette, & Parent, 2003; Wu et al., 2000). In addition, pallidostriatal inhibition of striatal interneurons can possibly disinhibit MSN activity. Finally, via local boutons of GP neurons (Fujiyama, Nakano, et al., 2015; Mallet et al., 2012), mutual inhibition (Bugaysen, Bar-Gad, & Korngreen, 2013; Mastro et al., 2017; Sadek et al., 2007) may work to terminate transient excitation of the GP.

In the case of M2 input to GP, the GP_STN_ pathway can also be activated, as well as the GP_CPu_ pathway described above (Fig. 10B). M2-GP_STN_ circuitry may be more sensitive to timing, because the GP and STN form bidirectional connections, and the cortico-pallidal pathway can act as fast as the hyperdirect pathway. These two pathways may compete: if GP_STN_ is activated first, it will suppress STN; if STN is activated first, GP will be excited directly by the cortex and via the STN, in turn inhibiting the STN. Taken together, the results suggest that the net effect of M2-GP_STN_ pathway activity is likely to be suppression of the STN. However, basal ganglia activity may be strongly affected by competition between M2-pallidal and hyperdirect pathways. Because the hyperdirect pathway contributes to cessation of ongoing actions, the cortico-pallidal pathway may cancel this cessation signal. Mallet et al. (2015) proposed a two-step model for cancellation via cooperation between the STN and GP, especially arkypallidal neurons. Our current findings shed light on the linking of complex stop/cancel sequences.

### Functional differentiation between M1 and M2

The overall effect of cortico-pallidal innervation may be suppression of the hyperdirect pathway, and its timing and efficacy related to the selective cortical innervation of GP cell-type(s). Therefore, our first assumption concerning the origin of the cortico-pallidal pathway should be revisited here. A paper reported that PT neurons issuing axon collaterals in the STN did not provide collaterals to the GP (T. Kita & Kita, 2012). If the hyperdirect and cortico-pallidal pathways originate from distinct neurons, the timing of activity will be substantially affected.

The findings of recent *in vivo* experiments in rodents are contradictory regarding the presence of functional differentiation between M1 and M2 during movement tasks, implying task-dependency (Makino, Hwang, Hedrick, & Komiyama, 2016). On one hand, M2 contributes to the preparatory function (Svoboda & Li, 2018), as observed in primates (Wise, 1985), or increases its influence on other cortical areas during the preparatory phase (Makino et al., 2017). PT neurons have been strongly implicated in preparatory processes as well as movement (Li et al., 2015). GP neurons are also suggested to compute action selection (Bogacz et al., 2016; Goldberg & Bergman, 2011), which likely relates to preparation of movement; and the M2-pallidal pathway may therefore contribute. It has been suggested that M2 is involved in the integration of movement with sensory or internal information (Barthas & Kwan, 2017; Saiki et al., 2014), including posture coding (Mimica, Dunn, Tombaz, Bojja, & Whitlock, 2018). It is possible that M2-GP innervation reflects sensory signal information, and contributes to the fine tuning of ongoing movement such as the adaptation of chosen behaviors. Along with possibly involving the dopamine system, the M2-GP pathway may relate more plastic and integrated phases of movement, which may require sensitive control using two GP cell types. Natural action is a highly complex system, which involves the cerebrum, brain stem, spinal cord, and midbrain nuclei (Arber & Costa, 2018; Bostan, Dum, & Strick, 2013; Kelly & Strick, 2004). Concurrent and similar activities of M1 and M2 neurons have been reported, at least for certain movements (Saiki et al., 2014; Soma et al., 2017). Parallel connections between M1 and M2 also affect the aforementioned neural circuitry (Ueta, Hirai, Otsuka, & Kawaguchi, 2013; Ueta, Otsuka, Morishima, Ushimaru, & Kawaguchi, 2013). To conclude, the functional relevance of cortico-pallidal projections, M1/M2, and Cg differentiation for both behavioral output and neural circuitry remain to be resolved in detail.

## Materials and methods

Animal experiments were approved and performed in accordance with the guidelines for the care and use of laboratory animals established by the Committee for Animal Care and Use and the Committee for Recombinant DNA Study of Doshisha University. All efforts were made to minimize animal suffering and the number of animals used.

### Animal surgery for injection of neural tracers and viral vectors

Wistar SLC rats (Japan SLC Inc., Hamamatsu, Japan; *N* = 65 of both sexes for electrophysiological experiments and *N* = 30 male rats for morphological experiments) were anesthetized with intramuscular injection of a mixture of ketamine (Ketalar; Daiichi-Sankyo, Tokyo, Japan; 40 mg/kg) and xylazine (Bayer HealthCare, Tokyo, Japan; 4 mg/kg). A small amount (0.05 mL) of the mixture was additionally injected every 15 min during any prolonged surgery (>1 h). Body temperature was monitored and controlled at 37°C with the aid of a heating device (World Precision Instruments [WPI], Sarasota, FL, USA). Craniotomy was performed with a drill at an appropriate position on the skull based on the rat brain atlas (Paxinos & Watson, 2007). A glass pipette (tip diameter, 30–60 µm) was used for all injections. For anterograde tracers, biotinylated dextran amine (BDA, 10 kDa; 10% solution dissolved in PB; Thermo Fisher Scientific, Waltham, MA, USA) was injected using either a brief air pulse (10–20 psi for 40–100 ms) controlled with a coordinated valve system (PV 820, WPI) or using electrophoresis (0.5–2 μA of positive current, 7 s ON/7 s OFF for 80 cycles) with a current controller (WPI). PHA-L (Vector Laboratories, Burlingame, CA, USA; 2.5% dissolved in 10 mM Na_2_HPO_4_, pH 8.0) was injected using electrophoresis (1–5 μA positive current, 7 s ON/7 s OFF for 80 cycles). To reduce number of rats, BDA and PHA-L were injected into different cortical areas in individual rats, and brain sections derived from each rat were used for quantitative morphological analysis in GP, CPu, and STN (Fig. 1, .7, Supplementary Fig. 1). AAV vectors encoding fluorophores were injected using air pressure (1.5 × 10^10^ vg [vector genomes]/µL, *AAVdj-hSyn-hChR2(H134R)-mCherry*; the plasmid a gift from Karl Deisseroth; Addgene plasmid #26976; http://n2t.net/addgene:26976; RRID: Addgene_26976). For retrograde tracers, cholera toxin subunit B (CTB) conjugated with Alexa Fluor 488 or 555 (Thermo Fisher Scientific) and fluorophore-labeled beads (Green Retrobeads IX, LumaFluor Inc., Durham, NC, USA; FluoSphere Orange 0.04 µm, Thermo Fisher Scientific) were injected using air pressure. For labeling cortical neurons, injections were performed at two to three depths typically at 400 μm-intervals. The stereotaxic coordinates (Paxinos & Watson, 2007) of injections in the centers of each cortical area were as follows (the actual injection locations were normalized based on skull size using the distance between bregma and lambda): for rostral M1, 2.0 mm rostral from bregma (A/P r2.0) and 2.5 mm lateral from the midline (M/L 2.5), depth 0.5–1.2 mm from cortical surface; for caudal M1: A/P 0.0, M/L 2.6, depth 0.5–1.2; for M2: A/P r4.2, M/L 1.9, depth 0.5–1.2 at a 30° angle rostral from vertical; for lateral orbitofrontal area (LO): A/P r3.4, M/L 2.9, depth 4.0. Because the Cg is located at the medial surface of the frontal cortex and is enclosed by M2, contamination of M2 following vertical injections into the Cg would have been unavoidable. Therefore, for injections into the Cg, the contralateral medial frontal area was removed using an aspiration needle and vacuum pump to expose the medial surface of the frontal cortex of the targeted hemisphere. An injection electrode was then inserted into the Cg at an angle of 45° (A/P r1.8, M/L 0.5) and the contents of the electrode ejected into the Cg at two depths (0.5 and 1.0) using air pressure. Tracer injections into the striatum were performed in two ways. To visualize striatum subregion-specific projections (Fig. 8), iontophoresis was applied at a single location at A/P r1.0, M/L 3.5, depth 4.2, for injection into CB-negative dorsolateral striatum, and at A/P 0.8 mm caudal to bregma (c0.8), M/L 4.0, depth 3.9 for CB-positive areas. For efficient retrograde labeling of GP neurons projecting to the dorsal striatum, pressure injection was applied in three tracks (two depths for each track) at A/P r2.0, M/L 2.5, depth 3.8; A/P r1.0, M/L 3.5, depth 4.2; and A/P 0.0, M/L 3.7, depth 4.0. Micro-iontophoretic injections into subregions of the GP were located at A/P c1.2 for the rostral GP, A/P c1.6 for the central GP, and A/P c2.0 for the caudal GP. For these injections, the M/L and depth positions were 3.2 and 6.2, respectively. Tracers were injected into the STN at A/P c3.5, M/L 2.5, and depth 7.5 using air pressure. After injections, the skin was sutured and 2.5 mg/kg of butorphanol (Vetorphale, Meiji Seika Pharma, Tokyo, Japan) was subcutaneously injected as an analgesic. The animals were allowed to recover before further experimentation. We allowed 2–4 d of survival time for BDA or retrograde tracers, 5–8 d for PHA-L, and 2–4 weeks for AAV. The age of rats ranged 7 to 12 weeks for morphological experiments.

### Immunohistochemistry

After the survival period, rats were deeply anesthetized with an intraperitoneal injection of sodium pentobarbital (100 mg/kg; Kyoritsu Seiyaku Corporation, Tokyo, Japan) and perfused with pre-fixative (sucrose 8.5% w/v, MgCl_2_ 5 mM; dissolved in 20 mM PB) followed by fixative (2 or 4% paraformaldehyde and 0.2% picric acid with or without 0.05% glutaraldehyde in 0.1 M PB, pH 7.4) through the cardiac artery. The brains were post-fixed *in situ* for 2–3 h, and then removed and washed with PB several times. Sagittal or coronal sections 50-µm thick were cut using a vibratome (Leica VT1000, Leica Instruments, Wetzlar, Germany) or freezing microtome (Leica SM 2000R), and stored in PB containing 0.02% NaN_3_ until further use.

For immunoreaction, sections were incubated with primary antibody diluted in incubation buffer consisting of 10% normal goat serum, 2% bovine serum albumin, and 0.5% Triton X in 0.05 M Tris buffered saline (TBS) overnight at room temperature (RT) or for 2–3 d at 4°C. For immunofluorescence, after rinsing with TBS three times, the sections were incubated with the secondary antibodies conjugated with fluorophores for 3 h at RT. After three rinses, the sections were dried on glass slides and coverslipped with antifade mounting medium (ProLong Gold, Vector). For brightfield specimens, the sections were incubated with a biotinylated secondary antibody followed by rinses and then reacted with ABC solution (1:200 dilution; Vector Elite) for 3 h at RT, and visualized with 3,3′-diaminobenzidine (DAB), Ni-DAB or Tris-aminophenyl-methane (TAPM). The sections were dehydrated with a graded series of ethanol, delipidated with xylene, and finally embedded with M·X (Matsunami Glass Ind., Ltd., Osaka, Japan). The primary and secondary antibodies used in this study are listed in Supplementary Tables 1 and 2.

### Image acquisition

Photomicrographs of brightfield specimens were captured using a CCD camera (DP-73, Olympus, Tokyo, JAPAN) equipped with a BX-53 microscope (Olympus) using 4× (numerical aperture [N.A.] 0.13), 10× (N.A. 0.3), 40× (N.A. 0.75), 60× (N.A. 0.9) or 100× (N.A. 1.4; oil-immersion) objectives. Photomicrographs were analyzed using Fiji (a distribution of Image J) (Schindelin et al., 2012), and Adobe photoshop (Adobe Systems Incorporated, Sam Jose, CA, USA). The brightness of digitized images was adjusted using the adjust-level function of these applications. To obtain a multifocus image, images were captured with 1-μm steps and processed with the “extended depth of focus” Fiji plugin. Fluorescent images were captured using an Orca Spark CMOS camera (Hamamatsu photonics, Hamamatsu, Japan) or a DP-73 camera equipped with a BX-53 microscope. To quantify the molecular expression patterns of GP neuron types, immunofluorescent images were acquired using a confocal microscope (FV1200, Olympus) with 40× (N.A. 0.95) or 100× (N.A. 1.35; silicon oil immersion) objectives.

### in vitro slice recordings

Basal ganglia neurons were recorded using *in vitro* whole cell patch clamp. Rats of both sexes (*N* = 65; postnatal 30–65 d) were deeply anesthetized with isoflurane and perfused with 25 mL of ice-cold modified artificial cerebrospinal fluid (ACSF; N-methyl-D-glucamine, 93; KCl, 2.5; NaH_2_PO_4_, 1.2; NaHCO_3_, 30; HEPES, 20; glucose, 25; sodium ascorbate, 5; thiourea, 2; sodium pyruvate, 3; MgCl_2_, 10; and CaCl_2_, 0.5; all in mM; pH was adjusted to 7.3 with HCl). All ACSFs were continuously aerated with 95/5% O_2_/CO_2_. Brains were removed and immersed in ice-cold modified ACSF for 2 min. Coronal slices 300-μm thick were cut using a vibratome (7000smz-2, Campden, Leicestershire, UK) and incubated with modified ACSF at 32°C for 15 min. The slices were transferred to normal ACSF (NaCl, 125; KCl, 2.5; CaCl_2_, 2.4; MgCl_2_, 1.2; NaHCO_3_, 25; glucose, 15; NaH_2_PO_4_, 1.25; pyruvic acid, 2; lactic acid, 4; all in mM) at RT. After 1 h of recovery, slices were moved into a recording chamber thermostatted at 30°C. A whole-cell glass pipette of 4–6 MΩ was filled with intracellular solution (K-gluconate, 130; KCl, 2; Na_2_ATP, 3; NaGTP, 0.3; MgCl_2_, 2; Na_4_EGTA, 0.6; HEPES, 10; biocytin, 20.1; all in mM). The pH was adjusted to 7.3 with KOH, and the osmolality was adjusted to ∼290 mOsm. Target brain regions were identified with the aid of a fluorescence microscope (BX-51WI, Olympus) using a ×40 water-immersed objective lens. Current clamp recordings were low-pass filtered at 10 kHz and recorded using EPC10 (HEKA Elektronik Dr. Schulze GmbH, Lambrecht/Pfalz, Germany) with a sampling rate of 20 kHz. The series resistance was examined by applying a brief voltage pulse of −10 mV for 10 ms and was confirmed to be less than 25 MΩ during recording. Shortly (less than 1 min) after achieving whole cell configuration, the firing responses to 1-s depolarizing current pulses (maximum intensity was 1000 pA, increasing in 50-pA steps) were recorded in current clamp mode. Passive membrane properties were monitored as responses to 1-s hyperpolarizing current pulses.

For photoactivation of ChR2, a 470-nm light-emitting diode (LED; BLS-LCS-0470-50-22, Mightex Systems, Pleasanton, CA, USA) was used at full field illumination through a 40× water immersion objective. Five-millisecond blue light pulses were applied at a maximum total power of ∼4 mW, at which neuronal responses were saturated. In some experiments, low concentrations of TTX, 1 μM, and 4-amino pyridine (100 μM) were added to the ACSF to isolate monosynaptic currents (Petreanu, Mao, Sternson, & Svoboda, 2009; Shu et al., 2007). CNQX (10 μM) and AP5 (50 μM) were applied to inhibit glutamatergic synaptic currents (*N* = 10), and SR95531 (gabazine; 20 μM) was applied (*N* = 10) to prevent GABA_A_ receptor-mediated synaptic currents. All pharmacological reagents were purchased from Tocris Bioscience (Bristol, UK).

Slices were then fixed with a mixture of 4% paraformaldehyde, 0.05% glutaraldehyde, and 0.2% picric acid in 0.1 M PB overnight at 4°C. Fixed slices were rinsed with PB (3 × 10 min), and then re-sectioned into 50-μm-thick sections. The sections were incubated with 1% H_2_O_2_ in 0.05 M TBS for 30 min at RT to deplete endogenous peroxidases, then rinsed with TBS three times. The sections were incubated with CF350-conjugated streptavidin for 2 h (1:3000; Biotium, Inc., Fremont, CA) for fluorescent investigation of biocytin filled neurons. For brightfield microscopy, the sections were incubated with ABC (1:200; Elite, Vector) overnight at 4°C. Biocytin-filled neurons were visualized for light microscopy with Ni-DAB using H_2_O_2_ at a final concentration of 0.01%. The sections were dried on a glass slide and coverslipped with EcoMount (Biocare Medical, LLC, Concord, CA).

### Data analysis

#### Quantification of corticostriatal axon density

To quantify the distribution of cortical axons in the striosome and matrix, regions of interest (ROI) were first set as square fields of approximately 10 × 10 pixels and located in a striosome component expressing MOR. The size of the ROI was adjusted according to the size of the striosome. The fluorescence intensity of each pixel in the ROI was measured, and the median was used as the representative intensity for that ROI. Next, an ROI of the same size was set in the matrix close to the striosome, and the median fluorescence intensity of the matrix ROI was obtained. Each striosome ROI was accompanied by one matrix ROI. During ROI placement, areas containing axon bundles of passage were excluded. To reduce the variance of the injection size and number of labeled neurons, the fluorescence intensity of the matrix ROI was normalized to that of the striosome ROI. Thus, if axons were equally distributed in the striosome and matrix, the normalized intensity would be 1.0. To quantify axon density in CB-positive striatum, images of CB immunofluorescence were binarized, and the number of pixels containing cortical axons were counted. The median value of pixel intensity was used for representing projection strength.

### Quantification of cortico-pallidal axon density

Many thick cortical axon bundles with strong fluorescence passed through the GP, which largely disrupted proportional relationships between fluorescence intensity and axon varicosity (or bouton) density. Therefore, we manually counted axonal varicosities in each ROI using brightfield microscopic images. ROIs of 233 × 173 μm were selected as the locations of densest axon terminals. Because cortical boutons were observed in the striatum and STN of the same animal, these were also counted, and the resultant bouton density was compared with those of the GP to decrease the effect of variability of tracer injections.

### Tracing and quantification of labeled cortical or pallidal axons in the STN

Axons in serial sagittal sections were manually traced on paper using a drawing tube equipped with a microscope (OPTIPHOT, Nikon Instech Co., Ltd, Tokyo, Japan) with a ×40 objective (N.A. 0.7). The tracings were scanned and saved as digital data. The resolution of a digitized image was 0.48 µm/px. For quantification using Fiji, the digitized data were converted into a binary image and the presence of axons was determined pixelwise. For comparisons of spatial axon distributions in the STN among cortical areas of origin, the STN was divided into 10 equal parts along the A/P axis or into six equal parts along the dorsoventral axis. In each division, the numbers of pixels with and without cortical axons were counted, and the proportion of axon-containing pixels to all pixels was calculated for each division. The proportion data was also normalized to the maximum value for each injection. The mean values of 2 M/L plane sections (M/L, 2.3 mm and 2.7 mm) are indicated in Fig. 7E (*N* = 3 rats for M1, *N* = 2 for M2, and *N* = 2 for LO).

### Analysis of electrophysiological data

Our analysis method has been previously described (Mizutani et al., 2017; Oh et al., 2017). Briefly, slice recording data were analyzed using Igor Pro 7 (Wave metrics Inc., Portland, OR) with the aid of the Neuromatic plugin (http://www.neuromatic.thinkrandom.com)(Rothman & Silver, 2018) and handmade procedures. Neurons in the striatum, basal nucleus of Meynert, and GP possessed distinct firing properties and were readily distinguished. In the GP, GABAergic projection neurons and putative cholinergic neurons were distinguishable by cell morphology and firing properties. Putative cholinergic neurons were excluded from the present data. The locations of recorded cells were visualized with biocytin and manually plotted on a digital image.

### Statistical comparisons

Averaged data are provided as mean ± standard deviation unless otherwise noted. Data comparison among more than two groups was performed using one-way ANOVA followed by a post hoc Tukey test, using R software (http://www.r-project.org/; R Project for Statistical Computing, Vienna, Austria). Data comparison between two groups was performed using the Wilcoxon rank sum test. For comparison of proportion values, Fisher’s exact test was used. For comparison of cumulative histograms, the Kolmogorov-Smirnov test was applied. Differences in data values were considered significant if *p* < 0.05. Significant differences are indicated using asterisks (*, *p* < 0.05; **, *p* < 0.01; ***, *p* < 0.001). All *p*-values are reported.

## Acknowledgement

The authors thank Dr. Yasuharu Hirai for discussion and comments. We also thank Dr. Yoon-Mi Oh for technical assistance.

## Figure-figure supplement legends

**Fig. 1-figure supplement 1.**
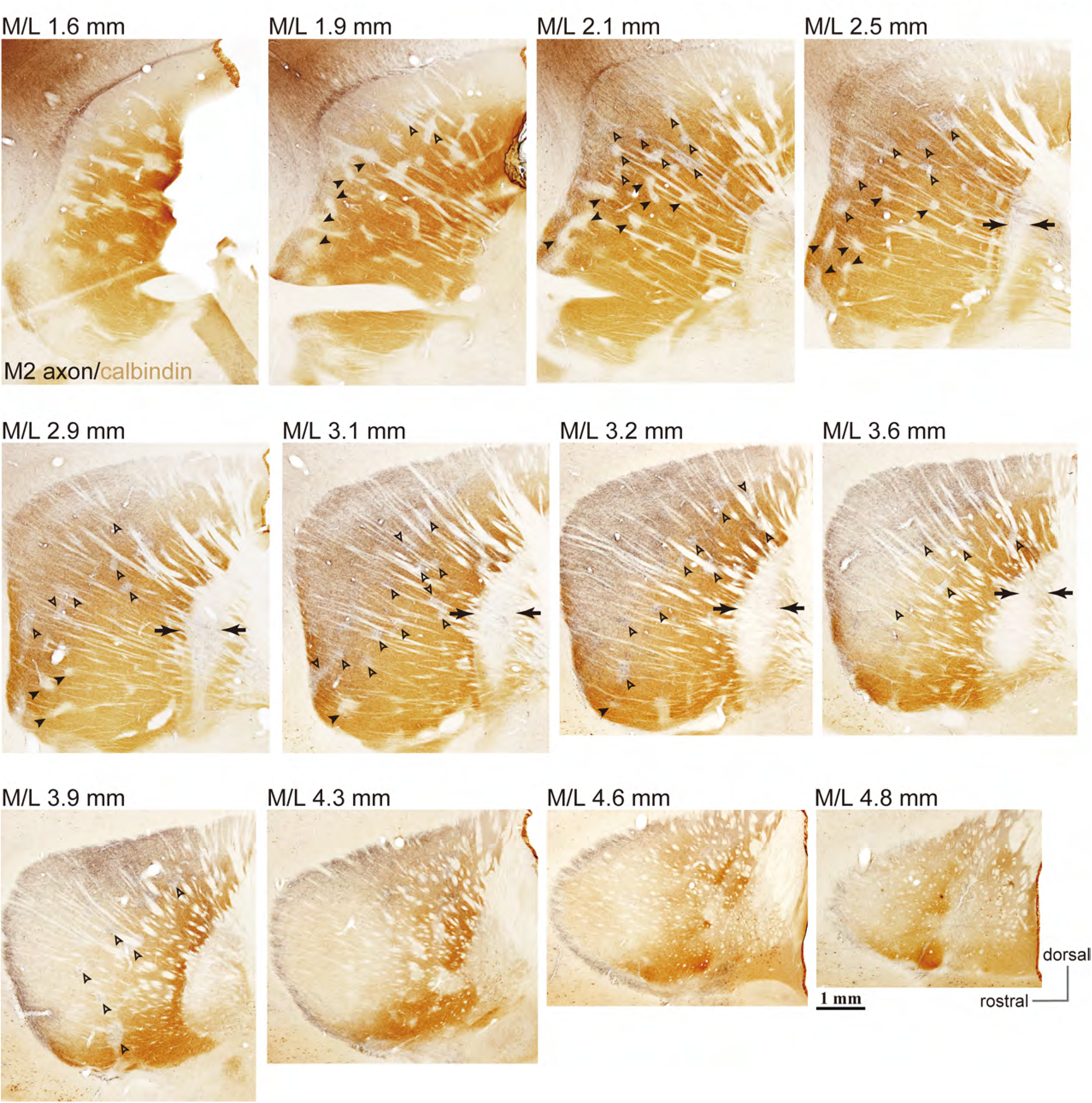
Example images of M2 projections to the striatum and GP (related to Fig. 1). Using BDA injections into M2, labeled axons are visualized in black and CB immunoactivity in brown. Axons are densely distributed in the striatum lateral to the midline (M/L 2.1–4.3 mm). In more lateral sections, axons were denser in CB(-) subregions, and were also found in striosomes (small CB[-] subregions) throughout the entire projection field (open arrowheads). In medial sections, M2 axons were observed in both CB(+) and CB(-) subregions. M2 axons were only faintly detected in ventral striosomes (filled arrowheads), whereas they were relatively dense in dorsal striosomes (open arrowheads). Note also the presence of M2 axons in CB(-) portions of the GP (arrows).

**Fig. 1-figure supplement 2.**
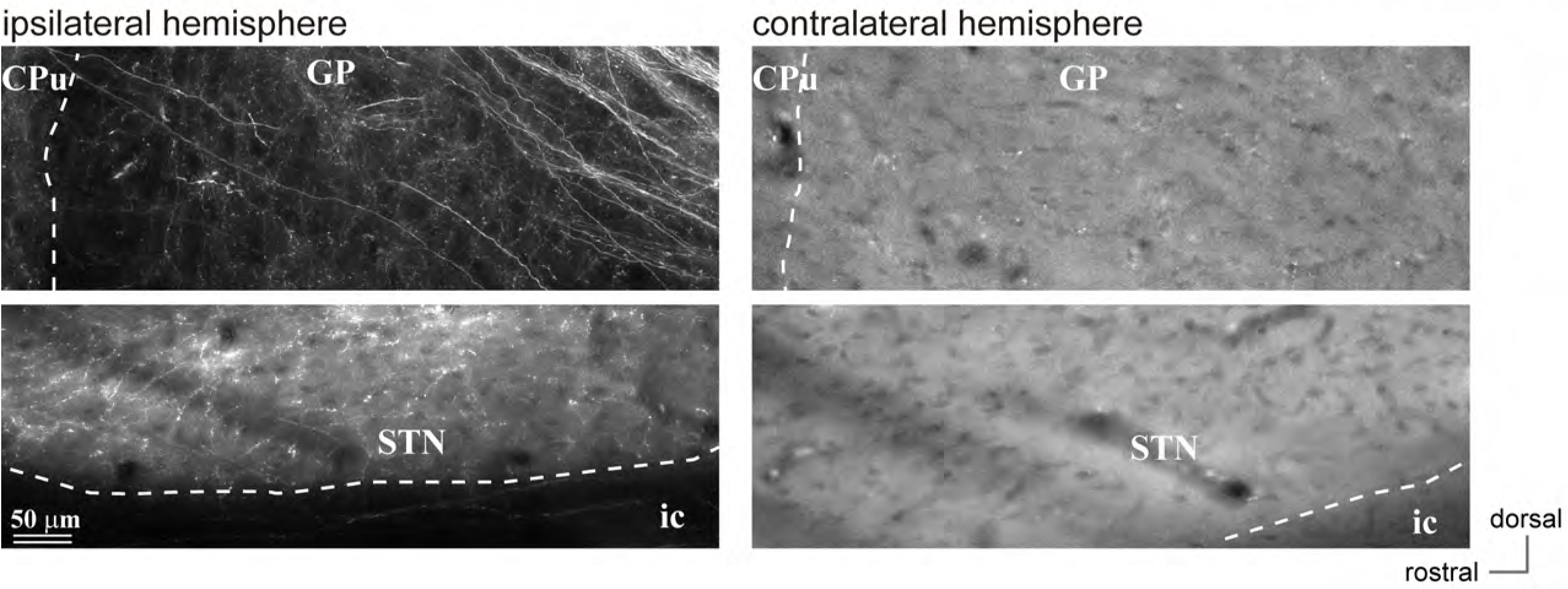
Cortico-pallidal projections are restricted to the ipsilateral hemisphere (related to Fig. 1). Representative images of AAV-labeled cortical axons in the basal ganglia originating in M2, in both hemispheres. **Left**, M2 axons were observed in the ipsilateral GP (**upper**) and STN (**lower**). In contrast, no fluorescent signal was observed in contralateral GP (**upper**) or STN (**lower**). The images in the contralateral hemisphere were captured with high exposures to better visualize brain structures, because no fluorescent signal was detected there. GP, globus pallidus; STN subthalamic nucleus; ic, internal capsule.

**Fig. 1-figure supplement 3.**
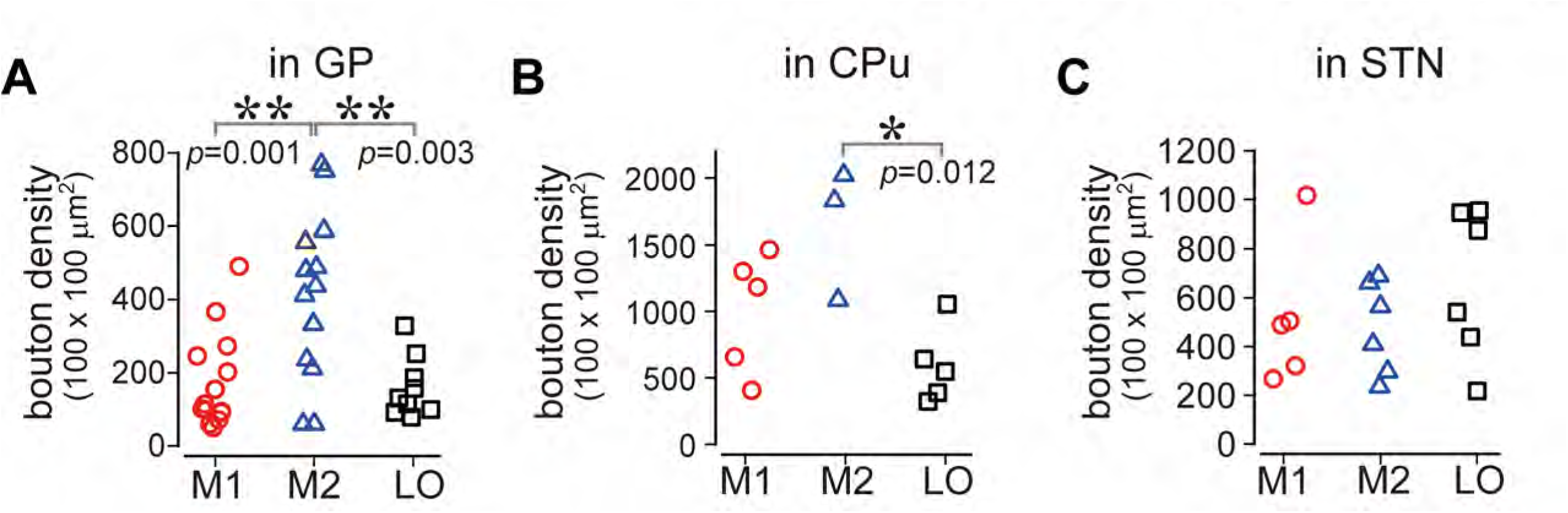
Number of cortical boutons in the striatum, STN, and GP originating from M1, M2, and LO (related to Fig. 1, also related to Fig. 5 and Supplementary Fig. 1).

**Fig. 7-figure supplement 1.**
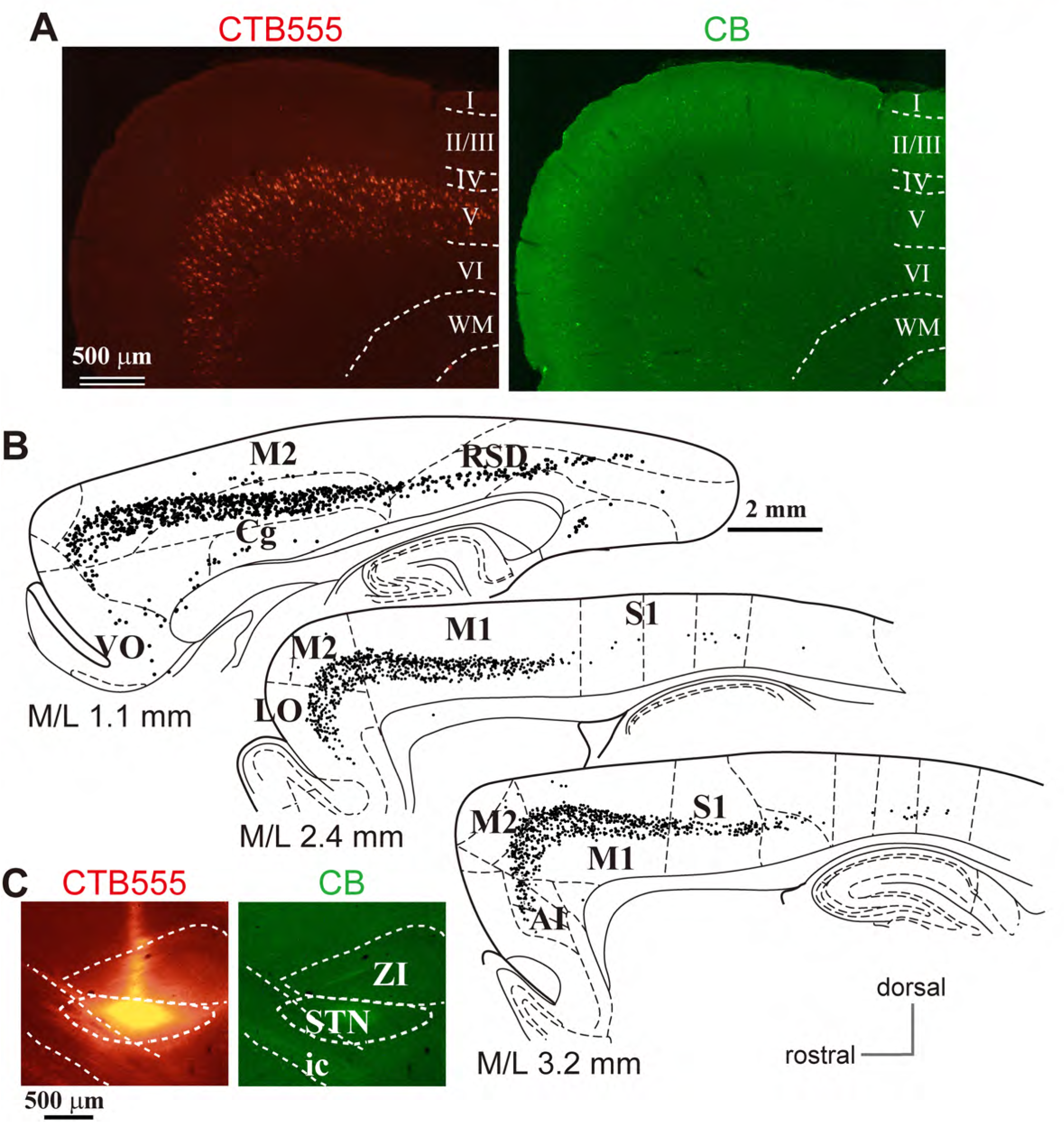
Representative distribution of STN-projecting cortical neurons (related to Fig. 5). (A) **Left**, retrogradely labeled cortical neurons projecting to the STN. Note that many L5A neurons were labeled in the frontal cortical area. **Right**, calbindin (CB) immunofluorescence (green) in the same section. Cortical layers are indicated with Roman numerals. (B) Plots of STN-projecting cortical neurons (black dots). Most labeled neurons are located in the frontal area including AI (the anterior insular area), the Cg (cingulate area), the LO (lateral orbitofrontal area), M1 (primary motor area), M2 (secondary motor area), and the VO (ventral orbitofrontal area). Some neurons are also located in S1 (primary somatosensory area) and the RSD (retrosplenial dysgranular area). (C) CTB555 was injected into the STN. ic, internal capsule; ZI, zona incerta. The effects of faint tracer leak to cortex and thalamus are considered negligible because very few labeled neurons are observed in L2/3 or L6, respectively.

## Supplementary materials

Orbitofrontal cortex (OFC) and anterior cingulate cortex (Cg) are involved in action (Schoenbaum et al., 2009). OFC provides behavioral flexibility via response inhibition and associative learning (but see previous reference). The OFC requires integration of sensory signals and internal states such as motivation with ongoing action selection and prediction of reward, to compute causal relationships between action selection and specific outcomes (Passingham & Wise, 2012). The Cg includes the rostral cingulate motor area (CMAr) (Morecraft & Van Hoesen, 1998), which lies in the cingulate sulcus, ventral to the presupplementary motor area, and in primates sends axonal projections to the spinal cord (Luppino, Matelli, Camarda, & Rizzolatti, 1994). The Cg and supplementary motor area (SMA) provide internal guides for action (Thaler, Chen, Nixon, Stern, & Passingham, 1995) correlated with motivation and voluntary behavior. The caudal cingulate motor area (CMAc), however, is preponderantly implicated in sensory-driven movement (Paus, 2001) and is thus related to action selection. Similarly to the OFC, the CMA promotes the switching of actions in response to changes in circumstances and internal desires (Amemori & Graybiel, 2012; Nakayama et al., 2015; Tanji, 1987), and is related to preparatory processes (Risterucci, Terramorsi, Nieoullon, & Amalric, 2003). CMAr projects to the M1, SMA, preSMA, premotor cortex, and brainstem, including the pons and spinal cord (Sessle & Wiesendanger, 1982). The OFC and Cg have different cognitive functions in primates and rodents (Bissonette et al., 2008; Bissonette et al., 2013; Bissonette & Roesch, 2015; Friedman et al., 2015; Nakayama et al., 2015; Sul et al., 2010). Thus, the OFC and Cg are important for motor behavior as well as being motor areas, and we therefore investigated axon projections to the basal ganglia from lateral orbitofrontal cortex (LO) and rostral Cg.

## Supplementary results

### Cortico-striatal and cortico-pallidal projections from LO and anterior cingulate cortex (Cg)

For comparison, we investigated axons from LO and Cg, which are also located in the rat frontal cortex LO axons were distributed in the medio-ventral part of the striatum, whereas Cg axons were distributed in the medial striatum (Suppl Fig. 1, 2). LO axons were sparse in the dorsal striatum but dense in the caudal and ventral part of the dorsal striatum as well as in the nucleus accumbens. In addition, these axons preferentially innervated the striosomes (Suppl. Fig. 1, arrowheads). In contrast, Cg axons mainly projected to the dorsomedial striatum with preference for the CB(+) subregions, namely the matrix (Suppl. Fig. 2). To a degree, the spatial distribution of Cg axons was similar to that of M2 axons; however, Cg axons did not preferentially project to the striosomes (Suppl. Fig. 2) (Averbeck, Lehman, Jacobson, & Haber, 2014; Friedman et al., 2015; Gabbott et al., 2005; Hintiryan et al., 2016).

LO axons also passed through the GP, but they branched less frequently, and axon collaterals were apparently not abundant (Suppl. Fig. 1). Actually, motor areas (both M1 and M2) provided more boutons than did LO, as expected based on the axon distribution. LO did not preferentially innervate the dorsal striatum or GP but preferred the nucleus accumbens, ventral pallidum, hypothalamus, and extended amygdala (Gabbott et al., 2005). LO projections to GP were not as dense as those to STN (Suppl. Fig. 1D1). The LO bouton density in the GP was approximately 30% of that in the striatum (Suppl. Fig. 1 D2). In contrast, the Cg projected to the medial GP and issued as many axon collaterals as did motor cortical areas (Suppl. Fig. 2).

## Supplementary Figure legends

**Supplementary Fig. 1.**
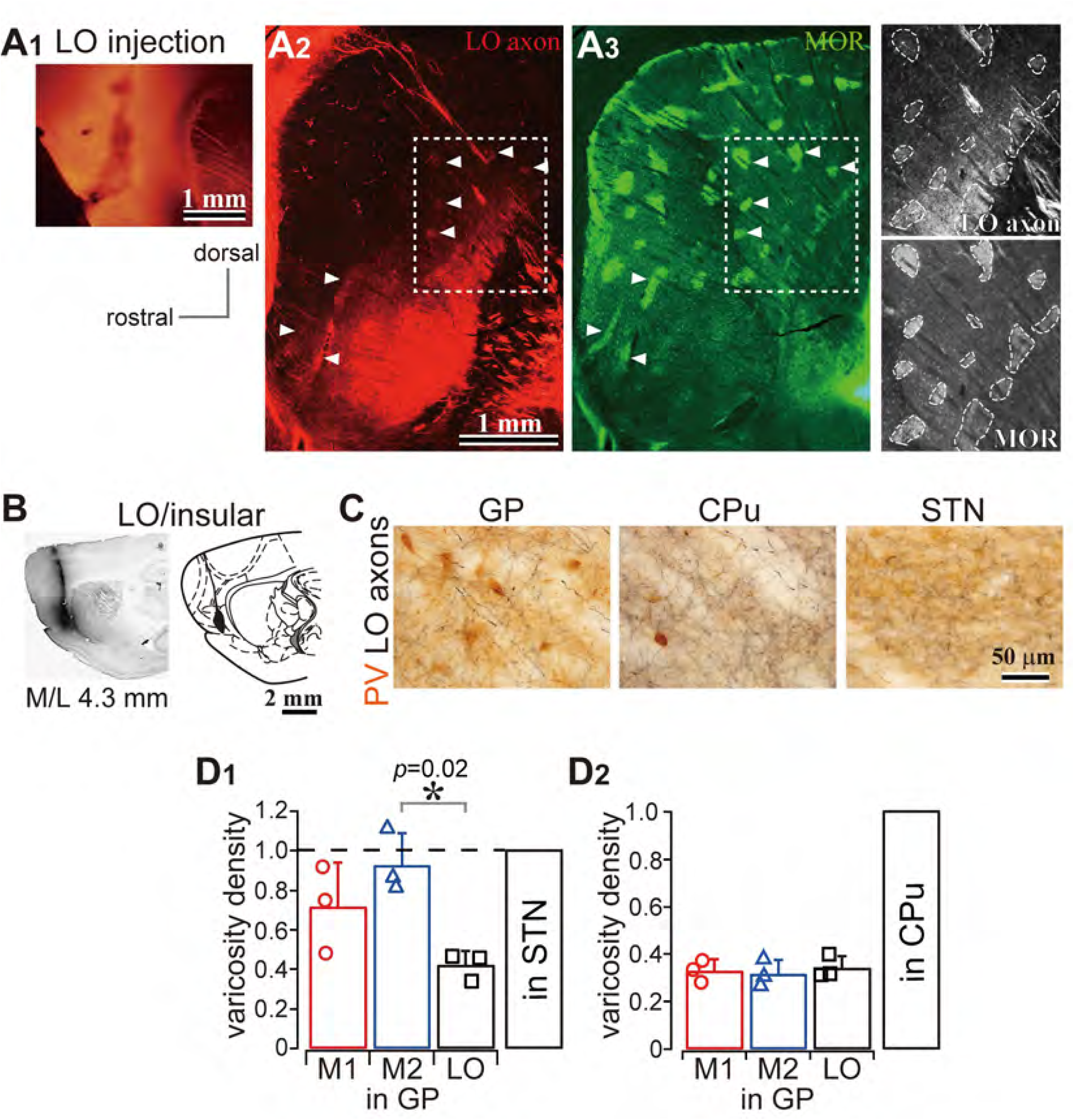
LO projections to the basal ganglia (A) LO projections to the striatum. (A1) AAV injections into the LO. (A2) A representative LO projection to the striatum. LO axons (red) are distributed in the caudal and ventral part of the dorsal striatum (M/L 2.6 mm). (A3) Immunofluorescence for μ-opioid receptor (MOR, green). Arrowheads indicate the striosomes, which are MOR(+), innervated by the LO axons seen in A2 and A3. Dotted square areas in A2 and A3 are magnified in the right-most column. (B) BDA injection into LO. (C) Brightfield images of LO axons in GP, striatum (CPu), and STN. Note that axon and terminal densities in GP were apparently lower than those in the striatum or STN. (E) Quantitative comparison of axon density in the GP normalized with that in the STN (D1) and striatum (D2). See also Fig. 1.

**Supplementary Fig. 2.**
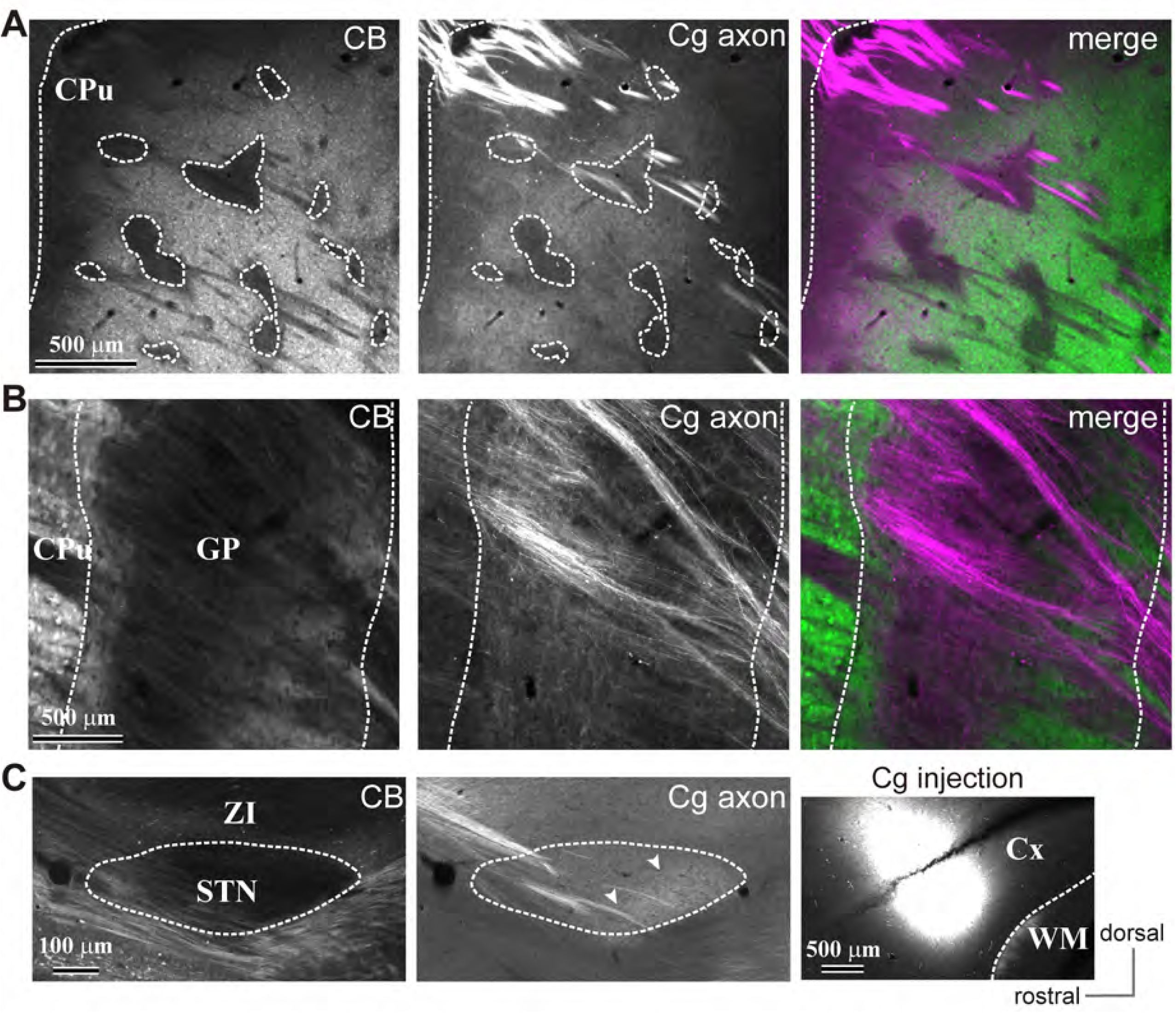
Cingulate cortex (Cg) projections to the basal ganglia (A) Cg projections to the striatum (CPu) in sagittal sections (M/L 2.10 mm). Cg axons are dense in the CB-negative subregions of the striatum. As in the case of M2 axons, Cg axons were less dense in the CB-negative striosomes. (B) Cg projections to the GP. Cg axon collaterals were dense in the CB(-) central GP. Thick axon bundles pass through the dorsal side of the medial GP (M/L 2.4 mm). (C) Cg projections to the STN. Cg axons are localized in the caudal part of the medial STN (arrowheads; M/L 2.4 mm).

**Suppl. Table 1.** Primary antibodies used in this study

| Antigen | Host<br>species | Supplier | Catalog number | RRID | Working<br>dilution |
| --- | --- | --- | --- | --- | --- |
| Calbindin | mouse | Sigma | C9848 | AB_476894 | 1:4000 |
| Calbindin | rabbit | Frontier<br>Institute | calbindin-Rb-Se-1 | AB_2571568 | 1:2000 |
| FoxP2 | rabbit | Abcam | ab16046 | AB_2107107 | 1:2000 |
| Lhx 6 | mouse | Santacruz | sc-271433 | AB_10649856 | 1:1000 |
| MOR | guinea<br>pig | Millipore | AB1774 | AB_91022 | 1:5000 |
| MOR | rabbit | Neuromics | RA10104 | AB_2156525 | 1:1000 |
| PHA-L | goat | Vector | AS2224 | AB_2315141 | 1:2000 |
| Parvalbumin | mouse | Sigma | P3171 | AB_2313693 | 1:4000 |
| Parvalbumin | guinea<br>pig | Synaptic<br>Systems | 195004/16 | AB_2156476 | 1:5000 |

MOR, μ-opioid receptor; PHA-L, *Phaseolus vulgaris* leucoagglutinin

**Suppl. Table 2.**
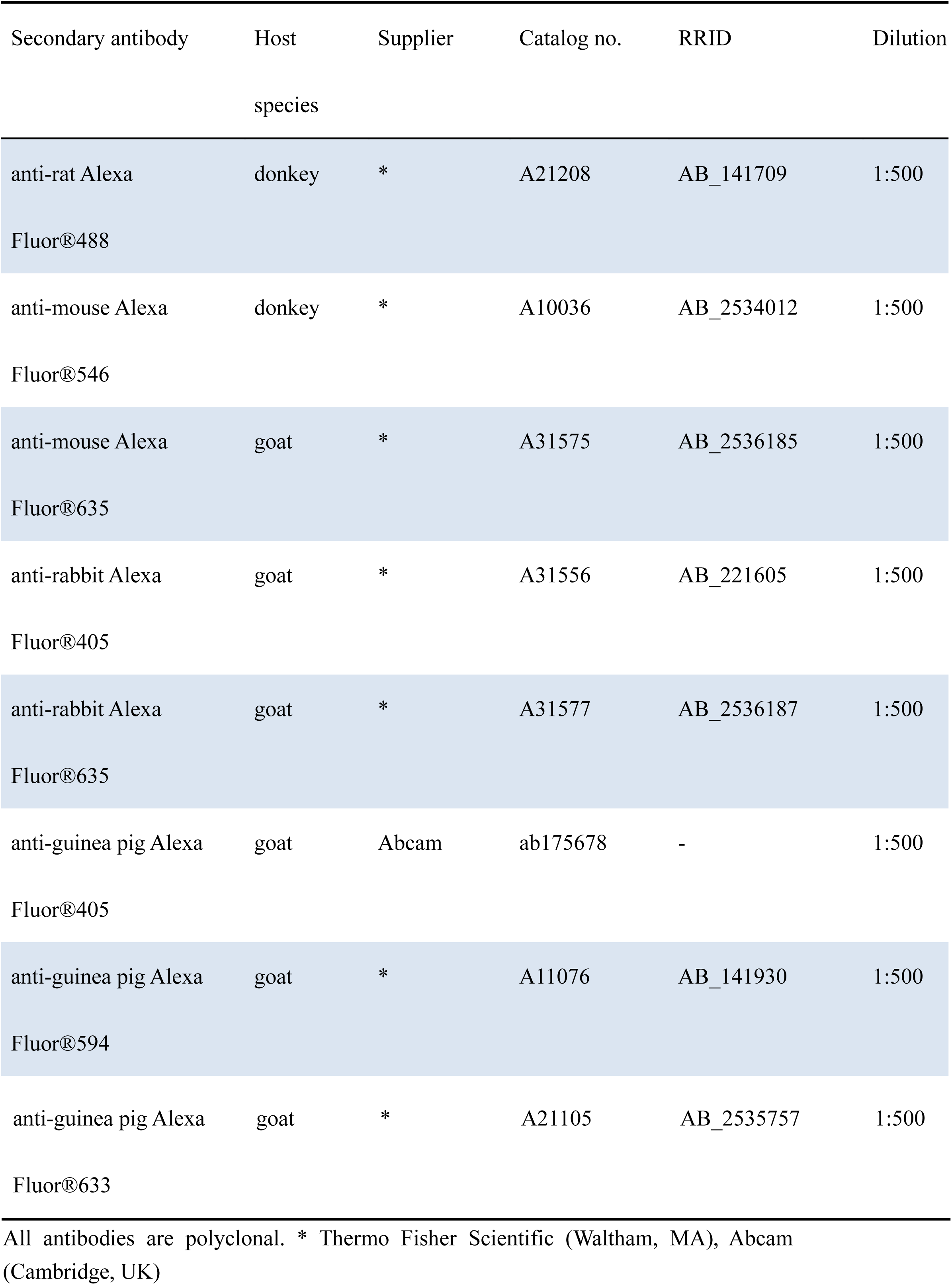
Secondary antibodies used in this study

